# L-lactic acid induces short- and long-term cardioprotective effects through MCT1 transport, induction of metabolic reprogramming, and gene expression modulation

**DOI:** 10.1101/2024.11.05.621926

**Authors:** Marina Cler, Soledad Pérez-Amodio, Laura Valls-Lacalle, Elena Martínez-Fraiz, Ignasi Barba, Freddy G. Ganse, Laura Nicastro, Cesare M. Terracciano, Antonio Rodríguez-Sinovas, Elisabeth Engel

## Abstract

Lactic acid is recognized as an alternative fuel source for various tissues and is acknowledged for its protective effects in the brain. However, its potential as a cardioprotective agent remains controversial. Here, we aimed to (1) evaluate the impact of acute L-lactic acid administration, given at the onset of reperfusion, on myocardial infarct size in isolated mouse hearts submitted to transient global ischemia, (2) assess the effects of chronic L-lactic acid exposure in living myocardial slices (LMS) from human hearts, and (3) elucidate the underlying mechanisms. Isolated mouse hearts were submitted to global ischaemia (35 min) followed by reperfusion (60 min), with L-lactic acid being or not administered during the first 15 min of reperfusion. L-lactic acid reduced infarct size by 23% at 20 mmol/L. An acidic Krebs induced less protection, and monocarboxylate transporter 1 (MCT1) inhibition with AR-C 141990 attenuated L-lactic acid’s protection to the level of acidic Krebs. ^1^H NMR spectroscopy revealed significant metabolic changes in L-lactic acid-treated hearts, with pathway enrichment analysis showing a nearly a 3-fold enrichment in pyruvate metabolism, fatty acid biosynthesis, and gluconeogenesis, suggesting a metabolic shift. Moreover, electrically stimulated human LMS treated with L-lactic acid for 48 h exhibited improved contractility and upregulation of structural and functional cardiomyocyte components, stemness-related markers, and pro-angiogenic proteins. These findings support a cardioprotective role for L-lactic acid in both short- and long-term contexts, mediated in part by its uptake through the MCT1 transporter, induction of metabolic reprogramming, and gene expression modulation.

## INTRODUCTION

Ischemic heart disease stands out as the most prevalent cardiovascular disorder and poses a significant economic challenge to public health systems worldwide. Current clinical guidelines from the European Society of Cardiology emphasize the urgent application of reperfusion therapies in patients with ST-segment elevation myocardial infarction (STEMI) ^1^. Nevertheless, given the magnitude of the problem and the fact that most patients end up with large infarct areas, there is growing interest in exploring new adjuvant therapies alongside reperfusion to improve patient prognosis. In this context, cardiac metabolism emerges not only as a key driver of myocardial infarction, but also as a potential target for intervention ^2^. Indeed, previous studies have provided evidence that the endogenous metabolite succinate plays a key role in ischemia-reperfusion injury, and that inhibition of succinate dehydrogenase with malonate reduces myocardial infarction in various animal models ^3–5^.

Lactate, a product of glucose metabolism, may hold significant importance in this context. Once considered merely as a dead-end product of anaerobic glycolysis, lactate is now recognized to play crucial roles in multiple cellular processes. Lactate serves as a substrate for oxidative phosphorylation, as a bridge connecting the glycolytic and the aerobic pathways, and as a major precursor for gluconeogenesis ^6^. It also regulates processes such as energy production, immune tolerance, wound healing, and cancer growth and metastasis ^6,7^. Notably, lactate is increasingly acknowledged as a protective compound in some tissues and under certain conditions. Experimental evidence suggests that lactate is neuroprotective after traumatic brain injury, in both animal models ^8^ and humans ^9^. In such scenarios, lactate has been proposed to serve as an alternative fuel source to optimize energy production and mitigate substrate deficiency ^9^. Similar protective effects have been described in rat hippocampal slices submitted to simulated ischemia-reperfusion and in a mouse model of middle cerebral artery occlusion, even when lactate was given immediately after ischemia^10–12^.

However, there have been relatively few studies analyzing the specific impact of lactate or lactic acid on the heart, particularly in the context of ischemic heart disease. It is known that elevated serum lactate levels are associated with increased infarct size and larger areas at risk ^13^. Additionally, lactate has vasodilator effects by stimulating NO release from the vascular endothelium ^14^, and acts as a free radical scavenger and antioxidant ^15^, action that can potentially influence final infarct size ^3^. Despite these findings there is a notable gap in the literature regarding the effects of lactate on myocardial infarction. To our knowledge, a single research groups has analyzed the impact of lactate or lactic acid on ischemia-reperfusion injury. In an in-situ rat model of transient coronary occlusion, Zhang and coworkers were not able to demonstrate any significant effect on infarct size of local, intramyocardial, administration of lactic acid when applied at the onset of reperfusion, despite it induced a reduction in pH in the right atrium and changes in the expression of signaling molecules ^16,17^. However, the intramyocardial route of administration is probably not the optimal to target reperfusion injury. Further, authors did not explore the mechanisms of protection and the relative contributions of tissue acidosis and transporter-dependent mechanisms. Finally, a single study has analyzed the effects of lactate in STEMI patients. Koyama and colleges employed a lactate Ringer’s solution during the reperfusion phases of an ischemic postconditioning protocol in patients with acute myocardial infarction. Postconditioning with lactate resulted in exceptional in-hospital clinical outcomes and during follow up ^18,19^. However, the study did not differentiate between the effects of postconditioning and lactate Ringer’s solution, nor did it compare results against a placebo group, complicating interpretation of the results. Therefore, the first aim of this study was to evaluate the impact of an acute administration of L-lactic acid, given at the onset of reperfusion, on myocardial infarct size in isolated mouse hearts submitted to transient global ischemia, and to elucidate the underlying mechanisms involved.

Further, lactate is increasingly considered as a promising bioactive molecule to boost cardiac endogenous repair using 3D mechanical metamaterial scaffolds ^20^. Lactate-releasing, cell-free, biomimetic scaffolds have been shown to reproduce tissue 3D organization and support neural migration, and when implanted in the postnatal mouse brain sustained neurogenesis and promoted long-term survival of newly generated neurons ^21^. Further, we have recently demonstrated that exogenous lactate promotes cardiomyocyte proliferation and cell cycle progression in mouse neonatal and human induced pluripotent stem cell-derived cardiomyocytes, with mouse cells responding to 20 mmol/L and human cells to 4-6 mmol/L ^22^. However, the effects of chronic administration of lactate in the adult myocardium are unknown. Therefore, the second aim of this work was to assess the functional effects of a chronic exposure to L-lactic acid in living myocardial slices from human hearts and to explore the mechanisms involved.

## METHODS

Animal studies complied with European legislation (Directive 2010/63/EU) on the protection of animals used for scientific purposes, with the Guide for the Care and Use of Laboratory Animals published by the US National Institutes of Health (NIH Publication No. 85-23, revised 1996, updated in 2011), and were approved by the Ethics Commettee of Vall d’Hebron Research Institute (CEEA31.15). The use of human samples adhered to the principles outlined in the Declaration of Helsinki and followed the European Guidelines for Good Clinical Practice. Human tissues from patients with heart failure and from two cardiac transplant donors whose organs were discarded for transplantation were provided by the NIHR Cardiovascular Biomedical Research Unit (Royal Brompton & Harefield NHS Foundation Trust and Imperial College London). The study received approval from the UK institutional ethics committee (NRES ethics number for biobank samples: 09/H0504/104 + 5; Biobank approval number: NP001-06-2015 and MED_CT_17_079) and Imperial College London. Informed consent was obtained from all patients or their families. A detailed description of the methods used in this study is available in the supplementary material online.

### Isolated, Langendorff-perfused, mice hearts

Adult male C57BL/6J mice (25-30g, 9-12 weeks) were used throughout the study. Animals were euthanized with an intraperitoneally overdose of sodium pentobarbital (1.5 g/Kg). Hearts were quickly removed and retrogradely perfused through the aorta with an oxygenated (95% O_2_: 5% CO_2_) Krebs solution (pH 7.4), using a constant flow Langendorff system, as previously described ^3^.

#### Concentration–response curves to L-lactic acid during normoxia

After a 30-minute equilibration period, four mice hearts were treated, under normoxic conditions, with Krebs buffer containing increasing concentrations of L-(+)-lactic acid (#199257, Sigma-Aldrich, USA), ranging from 1 to 50 mmol/L, each applied for 10 minutes and with pH adjusted to 7.4 (Fig. 1A). Osmolarity was maintained constant by addition of D-(+)-sucrose to match that of the highest L-lactic acid concentration. Lactate dehydrogenase (LDH) release was determined at the end of each concentration, and infarct size was measured at end of the experiment by TTC staining ^3^. Concentration-response curves for left ventricular developed pressure (LVdevP), left ventricular end diastolic pressure (LVEDP), heart rate, and perfusion pressure were fitted to sigmoid curves using the equation y=y0+a/[1+exp(-x-x0)/b], to determine the concentration causing half-maximal effect (EC50) and the maximal effect (Emax). LVdevP was defined as systolic pressure minus LVEDP.

**Figure 1.**
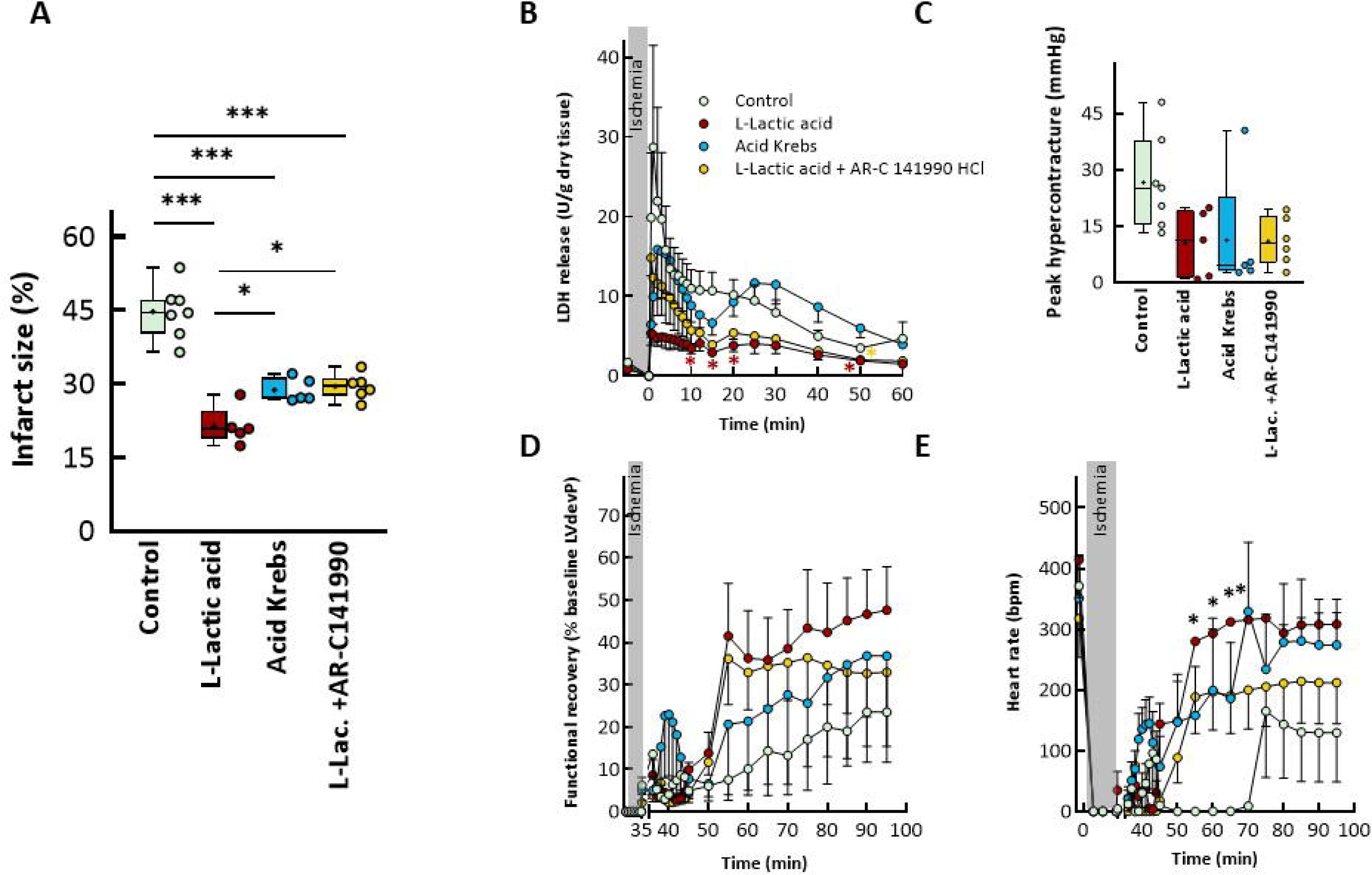
Effects of 20 mmol/L L-lactic acid (n=5), acid Krebs (n=5) and L-lactic acid + 1 mmol/L AR-C141990 (n=6), all given during the first 15 min of reperfusion, on (A) myocardial infarct size (* (p<0.05) and *** (p<0.001) indicate significant differences respect indicated groups (ANOVA and Tukey’s post hoc tests)), (B) LDH release (* (p<0.05) indicates significant differences between L-lactic acid or L-lactic acid+AR-C141990-treated hearts and controls (repeated measures ANOVA and Tukey’s post hoc tests)), (C) peak hypercontracture, (D) functional recovery and (E) heart rate (* (p<0.05) indicates significant differences between L-lactic acid-treated hearts and controls (repeated measures ANOVA and Tukey’s post hoc tests)) in isolated mice hearts submitted to transient global ischemia. Control hearts (n=7) were treated with standard Krebs buffer for the entire reperfusion. Data for infarct size and hypercontracture are shown as box plot depicting median (horizontal line), mean (+), and individual values (color symbols).

#### Effects of L-lactic acid on myocardial ischemia–reperfusion injury

Following a 30-minute stabilization period, mouse hearts were subjected to 35 minutes of global ischemia followed by 60 minutes of reperfusion (Fig. 2A-B), as described previously ^3^. Control hearts (n=7) received the standard Krebs buffer during the entire reperfusion (pH 7.4), whereas L-lactic acid-treated hearts (n=5/group) received Krebs containing L-(+)-lactic acid at 8 or 20 mmol/L, beginning at the onset of reperfusion and for the first 15 minutes. Selected concentrations were in the same order of magnitude as the EC50 found in the normoxic experiments.

**Figure 2.**
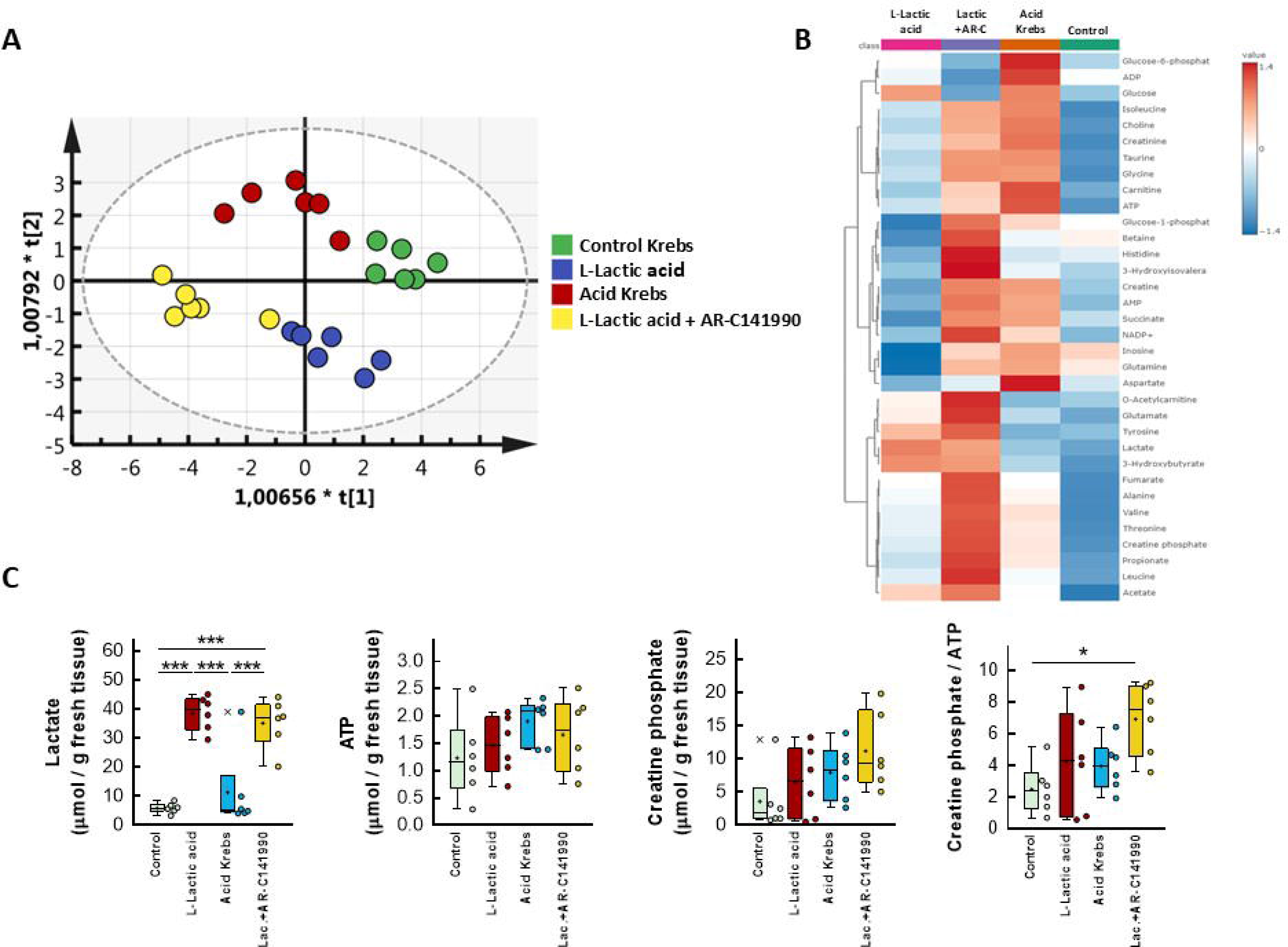
(A) Score plots corresponding to the discriminant analysis (OPLS-DA) of metabolites determined in myocardial samples from mice hearts submitted to 35 min global ischemia followed by reperfusion, and treated, during the first 15 min of reperfusion, with standard Krebs (n=7), or with Krebs containing L-lactic acid (n=5), with an acid Krebs (n=5) or with buffer added with L-lactic acid + 1 μmol/L AR-C141990 (n=6). (B) Heat-map showing changes in metabolites in all four treatment groups. (C) Quantification of lactate, ATP, creatine phosphate and the ratio creatine phosphate/ATP in hearts from the same four experimental groups. * (p<0.05) and *** (p<0.001) indicate significant differences between indicated groups (ANOVA and Tukey’s post hoc tests). Data are shown as box plot depicting median (horizontal line), mean (+), individual values (color symbols) and outlayers (x).

As pH of L-lactic acid-containing solutions was about 7.0, and to assess the relative contribution of acidosis to the observed effects, additional hearts were perfused with an acidic Krebs buffer (in mmol/L: NaCl 118, KCl 4.7, MgSO_4_ 1.2, CaCl_2_ 1.8, NaHCO_3_ 7.5, KH_2_PO_4_ 1.2, and glucose 11, pH 7.0) during the first 15 min of reperfusion (n=5) and results were compared with those obtained with 20 mmol/L L-lactic acid. In addition, to assess whether the effects of L-lactic acid were dependent on its transport into the cytoplasm, some hearts were simultaneously perfused with Krebs buffer containing 20 mmol/L L-(+)-lactic acid during the first 15 min of reperfusion while inhibiting its main transport system, the monocarboxylate transporter 1 (MCT1) ^7^, with 1 μmol/L AR-C141990 hydrochloride (#5658, Tocris, UK) (n=6). The concentration of the MCT inhibitor was selected based in previous publications ^5^.

Functional recovery (as a percentage of baseline LVdevP), heart rate, and hypercontracture (the difference between the maximum LVEDP during the initial minutes of reperfusion and the value at the end of ischaemia) were assessed in all cases.

#### LDH release and infarct size measurements

Cell death was indirectly quantified by measuring LDH release in samples from the coronary effluent by spectrophotometry, as previously described ^3^. Infarct size was determined in cardiac slices by 2,3,5-triphenyltetrazolium chloride staining at the end of reperfusion ^3^.

#### Analysis of myocardial metabolism

Cardiac metabolites were analyzed in 24 additional mouse hearts subjected to 35 minutes of global ischemia followed by 5 minutes of reperfusion, under the different treatments previously described (n=6/group) ^3^. OPLS-discriminant analysis and Fisher test were performed using SIMCA 14.0 (Umetrics, Sartorius, Germany). Metabolites were matched to metabolomics pathways using the Pathway Analysis and Enrichment Analysis features in Metaboanalyst 6.0.

### Human living myocardial slices (LMS)

Hearts from six patients with heart failure and from two donors who died from non-cardiovascular diseases were isolated, placed in cold cardioplegia solution, and stored in ice. Patient characteristics are detailed in supplementary table 1. Human LMS were quickly prepared as previously described ^23^, using a high-precision vibrating microtome (7000 smz-2; Campden Instruments, London, UK). Square cardiac slices (∼10x10mm) with homogeneous fiber alignment were cut and glued to rectangular custom-made, biocompatible polyethylene terephthalate (Taulman3D T-glase) 3D-printed holders, in a direction perpendicular to the fibers. LMS were placed in a sterile laminar flow cabinet and stretched to a physiological load (i.e., sarcomere length of 2.2 µm ^24^) by sterile, custom, stainless-steel stretchers. LMS slices were immediately cultured *in vitro* in custom sealed culture chambers, at 37°C, under continuous oxygenation (95% O_2_: 5% CO_2_). Culture medium consisted of medium 199 (Sigma-Aldrich, USA) supplemented with (in nmol/L) adrenaline 4, noradrenaline 4, dexamethasone 100, and 3,3L,5-Triiodo-L-thyronine 2.15, plus insulin-transferrin-selenium 0.1%, penicillin-streptomycin 2%, and ascorbic acid 20 μg/mL. LMS preparations were electrically stimulated using carbon electrodes at 0.5 Hz (10 ms pulse width, 15 V) and were randomly assigned to control or L-lactic acid-treated groups. L-(+)-lactic acid was directly dissolved in the cell culture medium at a concentration of 8 mmol/L and exposure lasted for 48h.

#### LMS contractility

Cardiac slices were affixed to a force transducer (HSE isometric force transducer F30 type 372, Harvard Apparatus, USA) using the 3D-printed holders, and stimulated at 0.5 Hz (10 ms pulse width, 20–30 V). The slices were stretched gradually until maximum isometric contraction was achieved. Force signals and peak amplitudes were recorded and analyzed using AxoScope and Clampfit software (Molecular Devices, San Jose, USA). Developed forces were normalized to cross-sectional area and reported as wall stress (mN/mm^2^).

#### Gene expression by quantitative real-time reverse transcription-polymerase chain reaction (qRT-PCR)

After culture with or without L-lactic acid, LMS were washed with PBS, detached from the holders, snap-frozen in liquid nitrogen, and stored at -20°C until analysis. Total RNA was extracted using the RNeasy® Mini Kit (Qiagen) according to the manufacturer’s instructions. RNA was retrotranscribed into cDNA with iScript™ cDNA Synthesis Kit (Bio-Rad Laboratories, USA). Gene expression was measured in triplicate with a QuantStudio 6 Pro Real-time PCR system (Applied Biosystems, USA) using iTaq™ Universal SYBR® Green Supermix (Bio-Rad Laboratories, USA) and pre-designed gene-specific primers (Supplementary Table 2). Relative mRNA levels were calculated by the 2^-ΔΔCt^ method using GAPDH as the housekeeping gene, and were expressed as fold change respect to non-treated LMS (without L-lactic acid).

#### Immunofluorescence analysis of lactate receptors and transporters and of angiogenic markers

After culture with and without L-lactic acid, LMS were washed with PBS, fixed in 4% formaldehyde solution (Thermo Scientific Scientific, Waltham, USA) for 15 min, and washed again in PBS. After fixation and blockade of non-specific binding, LMS were incubated overnight at 4°C with primary antibodies (mouse anti-CD31/PECAM-1, #sc-376764, Santa Cruc Biotechnology, dilution 1:50; rabbit anti-GPCR GPR81, #orb183872, Biorbyt, dilution 1:100; rabbit anti-SLC16A3 (MCT4), #orb6971, Biorbyt, dilution 1:100; or mouse anti-Von Willebrand factor, #sc-53466, Santa Cruz Biotechnology, dilution 1:100). Samples were then washed and incubated for 2h (RT) with the corresponding goat secondary anti-mouse or anti-rabbit antibodies marked with Alexa Fluor 488 or 568. Nuclei were counterstained with 4L,6-diamidino-2-phenylindole (DAPI) (#D9542, Sigma-Aldrich, USA, dilution 1:500) for 15 min. Immunolabelled samples were imaged using a wide-field epifluorescence microscope (Leica Thunder 3D Live Cell, Leica Biosystems Nussloch GmbH, Germany). Quantitative image analysis was performed using Fiji software. A minimum of 6 random images per sample were acquired and analyzed, and the result averaged.

#### Statistical analysis

Results are shown as mean ± standard error (SEM). Statistical differences were determined using GraphPad Prism 9.4.0. For two group comparisons, a two-tailed Student’s T-test was used. For multiple comparisons, one-way ANOVA followed by *post hoc* Tukey’s test (infarct size, cumulative LDH release, hypercontracture, metabolites) or two-way ANOVA followed by *post hoc* Bonferroni test (Frank-Starling relationship curves) were performed. Repeated measures ANOVA (MANOVA) followed by *post hoc* Tukey’s test was used for analysis of one intergroup and one intragroup variable (changes in LVdevP, heart rate, LVEDP, perfusion pressure). Differences were considered significant when p<0.05.

## RESULTS

### Impact of acute administration of L-lactic acid in isolated mouse hearts

#### Concentration–response curves to L-(+)-lactic acid during normoxia

Administration of L-(+)-lactic acid resulted in a concentration-dependent decrease in LVdevP. The data fitted well to sigmoid curves (Supplementary Fig. 1), showing strong correlation coefficients, with a half-maximal effective concentration (EC50) of 11.74 ± 6.10 mmol/L and a maximum effect (Emax) of 43.27 ± 13.87 % of baseline value. No significant changes were observed in left ventricular end-diastolic pressure (LVEDP), perfusion pressure, or heart rate at any of the tested concentrations. Additionally, no signs of injury were detected in any case, as indicated by the absence of LDH release and negligible infarct sizes at the end of the experiment.

#### Effects of L-lactic acid against ischemia-reperfusion injury in isolated mouse hearts

Administration of 20 mmol/L of L-lactic acid at the onset of reperfusion reduced the extent of infarction by about 23% in isolated mouse hearts submitted to 35 min of global ischemia (Fig. 1, Supplementary Fig. 2A-C), whereas the lower concentration of 8 mmol/L induced only a modest and non-significant effect. This protective action was associated with a significant reduction in LDH during reperfusion (Fig. 1, Supplementary Fig. 2D) and with a significant improvement in functional recovery (Fig. 1, Supplementary Fig. 2E). It is important to note that the addition of L-Lactic acid reduced the pH of the perfusate in a concentration-specific manner, reaching, after oxygenation, a pH of 6.97±0.05 when 20 mmol/L were used, as compared with that of control Krebs (7.65±0.02).

#### Contribution to acidosis and MCT1 to the cardioprotective effects of exogenous L-lactic acid

As a reduction in intracellular pH is known to exert a major cardioprotective effect ^25,26^, we initially compared the effects of 20 mmol/L L-lactic acid with those obtained with a perfusion, during initial reperfusion, with an acidic Krebs buffer containing a reduced amount of NaHCO_3_. Following oxygenation, the pH of this buffer was 7.01±0.06, similar to that of Krebs containing 20 mmol/L L-lactic acid. As shown in figure 1A, perfusion with this acidic Krebs significantly reduced infarct size compared to control hearts. This effect associated with a lower LDH release during reperfusion (Fig. 1B), a trend towards reduced peak hypercontracture (Fig. 1C), and with improved functional recovery (Fig. 1D-E). However, the cardioprotective effect of L-lactic acid was still significantly superior to that of the acidic Krebs in terms of infarct size reduction and LDH release (Fig. 1).

To determine whether the difference in protection between L-lactic acid and the acidic Krebs could be attributed to import of L-lactic acid into the cytoplasm of cardiomyocytes, we treated additional hearts with Krebs buffer containing both 20 mmol/L L-lactic acid and the MCT-1 inhibitor AR-C 141990. As depicted in figure 1, inhibition of the monocarboxylate transporter attenuated the cardioprotection induced by L-lactic acid alone, to a level similar to that found with the acidic Krebs.

#### Myocardial metabolic profile of mice hearts treated with L-lactic acid or acidic Krebs during reperfusion

Supervised classification of ^1^H NMR spectra obtained from myocardial extracts of isolated mice hearts treated with the different perfusates was able to discriminate between the different treatment groups (Fisher test: p< 0.0001, Q^2^ = 0.328) (Fig. 2A). These results suggest that hearts treated with the different perfusates exhibit distinct cardiac metabolic profiles.

Seventeen out of 37 metabolites were significantly different among the treatment groups (Figs. 2 and 3, Table 1). Hearts treated with a Krebs buffer containing L-lactic acid during the initial minutes of reperfusion showed significant increases in acetate and lactate, whereas those perfused with an acidic Krebs solution had enhanced concentrations of acetate, aspartate, and glycine (Table 1, Fig. 3). On the other hand, hearts treated with the MCT1 inhibitor AR-C 141990 simultaneously with L-lactic acid exhibited the highest number of altered metabolites. These included increased concentrations of acetate, glucose-1-phosphate, glycine, lactate, leucine, NADP^+^, O-acetylcarnitine, propionate, threonine, tyrosine, and valine, together with the creatine phosphate/ATP ratio (Table 1, Figs. 2 and 3).

**Figure 3.**
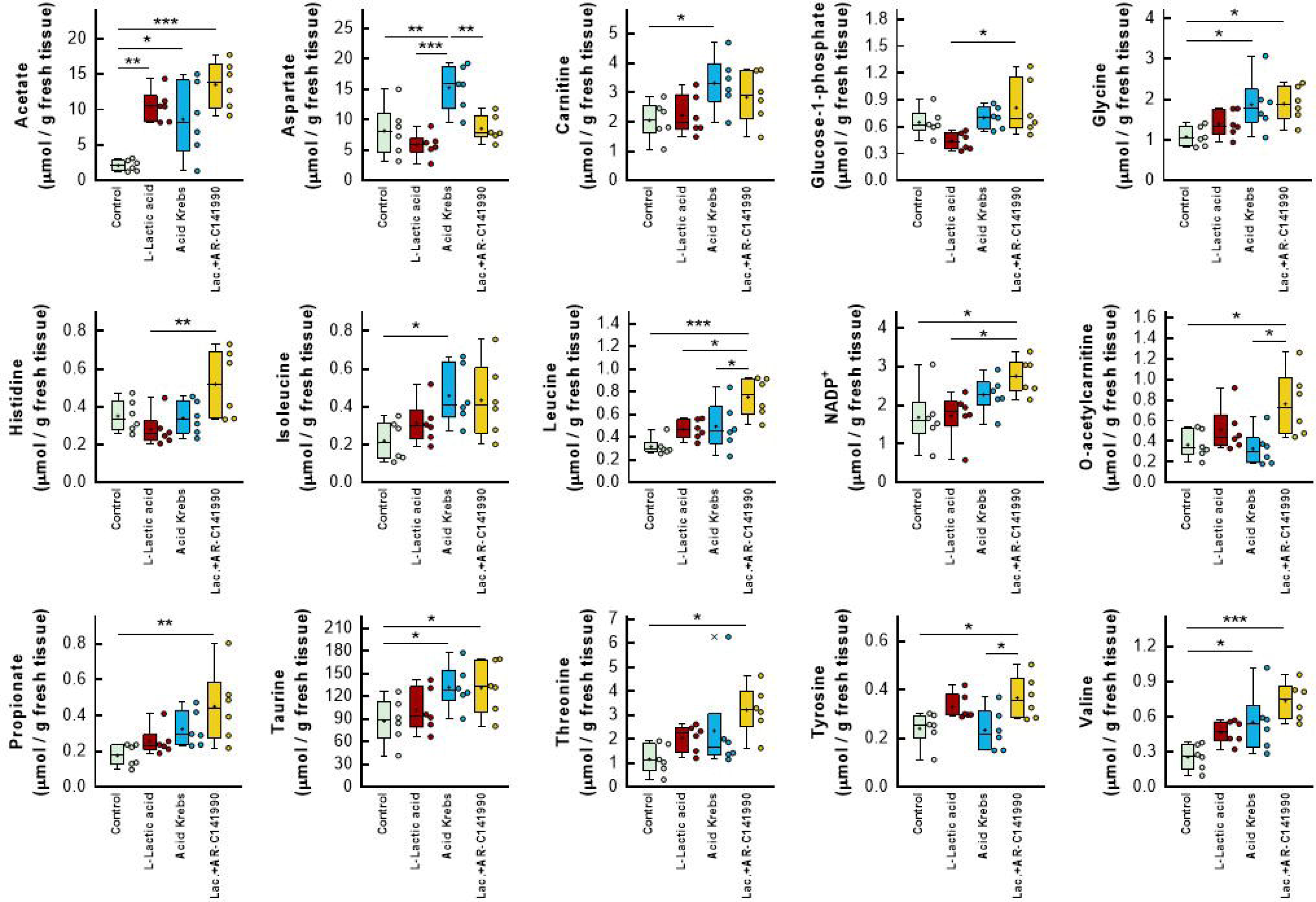
Quantification of metabolites by ^1^H NMR spectroscopy in samples from mice hearts submitted to 35 min global ischemia followed by reperfusion, and treated, during the first 15 min of reperfusion, with standard Krebs (n=7), or with Krebs containing L-lactic acid (n=5), with an acid Krebs (n=5) or with buffer added with L-lactic acid + 1 μmol/L AR-C141990 (n=6). * (p<0.05), ** (p<0.01) and *** (p<0.001) indicate significant differences between indicated groups (ANOVA and Tukey’s post hoc tests). Data are shown as box plot depicting median (horizontal line), mean (+), individual values (color symbols) and outlayers (x).

**Table 1.** Quantification of myocardial metabolites (mmol/g fresh tissue) analysed by ^1^H NMR spectroscopy. The tissue extracts employed for this analysis were obtained from hearts submitted to ischaemia (35 min) and reperfusion (5 min) (IR) with the different solutions. (n=6).

| Metabolite | Control | L-Lactic acid | Acid Krebs | L-Lactic acid + MCT-1 inhibitor |
| --- | --- | --- | --- | --- |
| 3-Hydroxybutyrate | 0.24 ± 0.05 | 0.66 ± 0.26 | 0.37 ± 0.07 | 0.65 ± 0.14 |
| 3-Hydroxyisovalerate | 0.3 ± 0.03 | 0.3 ± 0.01 | 0.33 ± 0.03 | 0.44 ± 0.07 |
| ADP | 0.59 ± 0.04 | 0.59 ± 0.12 | 0.62 ± 0.07 | 0.57 ± 0.08 |
| AMP | 1.04 ± 0.24 | 1.01 ± 0.25 | 1.44 ± 0.32 | 1.52 ± 0.3 |
| ATP | 1.23 ± 0.31 | 1.46 ± 0.22 | 1.9 ± 0.17 | 1.66 ± 0.28 |
| Acetate | 2.14 ± 0.32 | 10.51 ± 0.92 ** | 8.63 ± 2.15 * | 13.53 ± 1.35 *** |
| Alanine | 5.21 ± 0.99 | 8.19 ± 0.93 | 8.62 ± 2.39 | 11.52 ± 1.77 |
| Aspartate | 8.2 ± 1.69 | 5.85 ± 0.82 ## | 15.27 ± 1.52 ** | 8.57 ± 0.89 ### |
| Betaine | 0.68 ± 0.14 | 0.37 ± 0.07 | 0.62 ± 0.05 | 0.94 ± 0.29 |
| Carnitine | 2.05 ± 0.26 | 2.22 ± 0.28 | 3.32 ± 0.37 | 2.84 ± 0.36 |
| Choline | 0.68 ± 0.05 | 0.72 ± 0.07 | 0.83 ± 0.04 | 0.81 ± 0.12 |
| Creatine | 29.17 ± 3.88 | 27.68 ± 4.89 | 35.28 ± 4.57 | 35.92 ± 4.73 |
| Creatine phosphate | 3.6 ± 1.91 | 6.51 ± 2.16 | 7.95 ± 1.73 | 11.21 ± 2.37 |
| Creatinine | 0.91 ± 0.1 | 1.17 ± 0.07 | 1.56 ± 0.22 | 1.42 ± 0.43 |
| Fumarate | 0.14 ± 0.02 | 0.2 ± 0.03 | 0.2 ± 0.04 | 0.26 ± 0.04 |
| Glucose | 26.02 ± 1.94 | 31.26 ± 2.96 | 31.79 ± 6.47 | 24.93 ± 2.37 |
| Glucose-1-phosphate | 0.65 ± 0.06 | 0.44 ± 0.04 \$ | 0.7 ± 0.05 | 0.81 ± 0.13 |
| Glucose-6-phosphate | 2.81 ± 0.26 | 3.12 ± 0.31 | 3.88 ± 0.69 | 2.62 ± 0.53 |
| Glutamate | 4.65 ± 0.58 | 6.19 ± 0.52 | 5.35 ± 1.15 | 7.83 ± 0.85 |
| Glutamine | 5.45 ± 1.66 | 2.97 ± 0.41 | 6.3 ± 1.96 | 6.05 ± 1.31 |
| Glycine | 1.08 ± 0.1 | 1.4 ± 0.13 | 1.88 ± 0.28 * | 1.89 ± 0.18 * |
| Histidine | 0.35 ± 0.03 | 0.28 ± 0.04 \$\$ | 0.34 ± 0.04 | 0.52 ± 0.07 |
| Inosine | 2.21 ± 0.27 | 0.97 ± 0.14 | 2.48 ± 0.66 | 2.2 ± 0.46 |
| Isoleucine | 0.22 ± 0.04 | 0.32 ± 0.05 | 0.46 ± 0.06 | 0.44 ± 0.08 |
| Lactate | 5.96 ± 0.71 | 38.6 ± 2.45 *** , ### | 11.34 ± 5.62 | 35.22 ± 3.47 *** , ### |
| Leucine | 0.32 ± 0.03 | 0.47 ± 0.04 \$ | 0.5 ± 0.09 \$ | 0.75 ± 0.07 *** |
| NADP<sup>+</sup> | 1.69 ± 0.31 | 1.73 ± 0.25 \$ | 2.27 ± 0.19 | 2.75 ± 0.19 * |
| O-Acetylcarnitine | 0.37 ± 0.06 | 0.51 ± 0.09 | 0.33 ± 0.07 \$ | 0.76 ± 0.13 * |
| Propionate | 0.18 ± 0.03 | 0.26 ± 0.03 | 0.32 ± 0.04 | 0.45 ± 0.08 ** |
| Succinate | 0.65 ± 0.12 | 0.44 ± 0.07 | 0.98 ± 0.42 | 1.01 ± 0.31 |
| Taurine | 87.36 ± 12.17 | 101.68 ± 11.77 | 132.1 ± 11.82 | 131.14 ± 14.31 |
| Threonine | 1.17 ± 0.25 | 2.04 ± 0.23 | 2.35 ± 0.79 | 3.22 ± 0.41 * |
| Tyrosine | 0.24 ± 0.03 | 0.33 ± 0.02 | 0.23 ± 0.03 | 0.37 ± 0.04 * , # |
| Valine | 0.26 ± 0.04 | 0.47 ± 0.04 | 0.55 ± 0.11 * | 0.74 ± 0.06 *** |

\* Indicates a significant difference compared with hearts perfused with control Krebs. # Indicates a significant difference compared with hearts perfused with acid Krebs. \$ Indicates a significant difference compared with hearts perfused with 20 mM L-Lactic acid + MCT-1 inhibitor.

The heat-map shown in figure 4A shows changes in metabolites between individual cases in control and L-lactic acid-treated hearts. Pathway analysis identified multiple altered pathways in L-lactic acid-treated hearts as compared with control animals, including pyruvate metabolism, glycolysis/gluconeogenesis, and glyoxylate and dicarboxylate metabolism (Fig. 4B). Pathway enrichment analysis revealed almost a 3-fold enrichment in pyruvate metabolism, fatty acid biosynthesis, gluconeogenesis, amino sugar metabolism and vitamin K and porphyrin metabolisms, among others (Fig. 4C). Comparison of hearts perfused with acidic Krebs vs. controls resulted, both at the pathway and enrichment analysis, in lower p values than those found with L-lactic acid. Acid Krebs shared with L-lactic acid, at the pathway enrichment analysis, an almost 3-fold enrichment in fatty acid biosynthesis and vitamin K and porphyrin metabolisms, but differed in taurine and hypotaurine metabolism, phospholipid biosynthesis and carnitine synthesis (Supplementary Fig. 3). Notably, hearts treated simultaneously with L-lactic acid and AR-C 141990 shared with those treated with L-lactic acid alone 16 out of the top 25 metabolic pathways in the enrichment analysis (Supplementary Figure 4).

**Figure 4.**
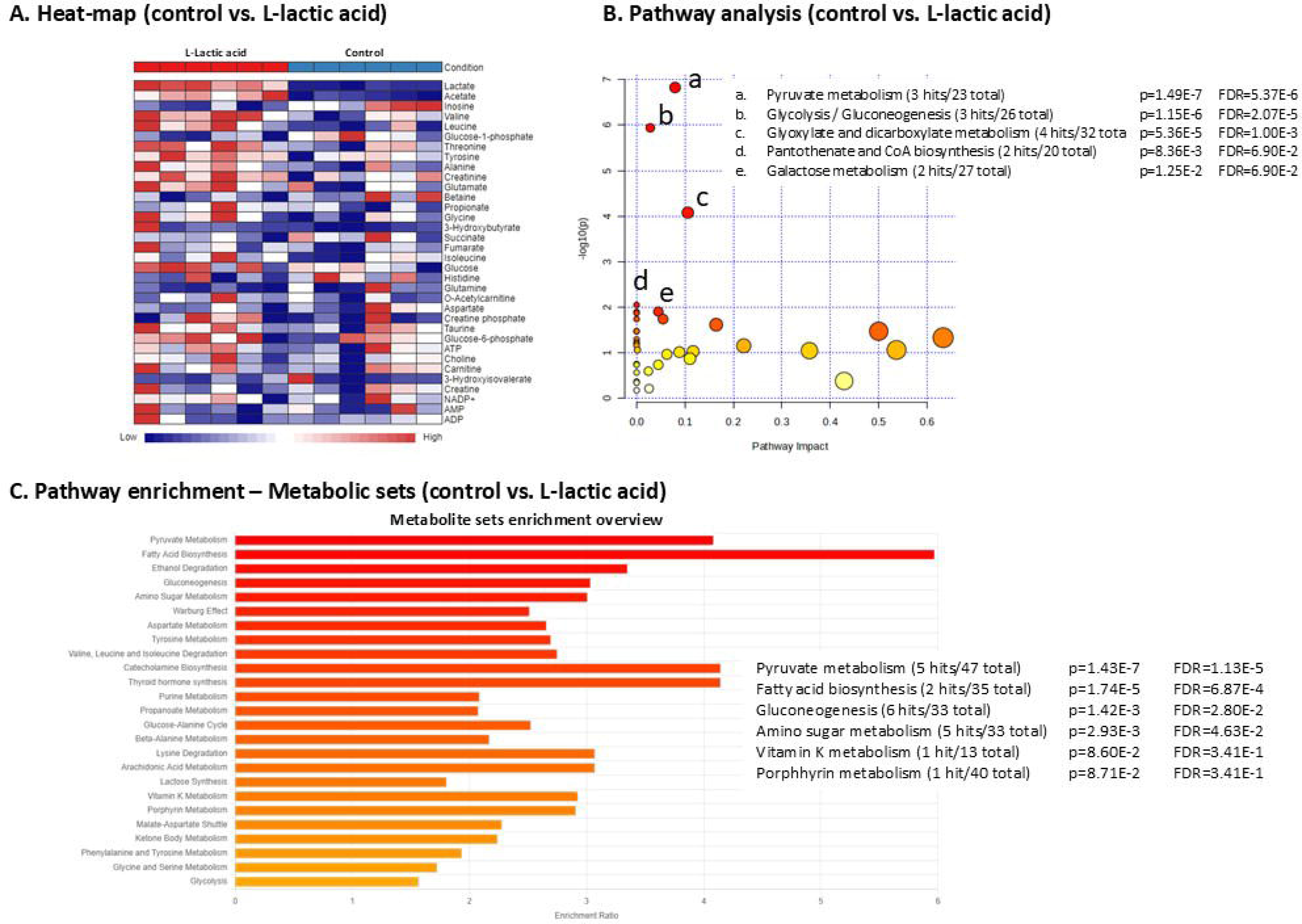
Non-targeted metabolomic analysis of hearts from mice hearts submitted to 35 min global ischemia followed by reperfusion, and treated, during the first 15 min of reperfusion, with standard Krebs (n=7), or with Krebs containing L-lactic acid (n=5). (A) Heat-map of metabolites between individual cases in control and L-lactic acid-treated hearts. (B) Pathway analysis of analyzed metabolites. (C) Pathway enrichment analysis of analyzed metabolites using metabolic datasets.

### Effects of a chronic exposure to L-lactic acid in LMS from failing human hearts

#### LMS contractility

Exogenous L-lactic acid significantly enhanced contractility of LMS from human failing hearts (Fig. 5A). Specifically, their maximum active force increased by more than 2-fold after treatment, reaching contraction values of 9.00 ± 0.75 mN/mm^2^ in L-lactic acid-treated LMS *vs.* 3.99 ± 0.59 mN/mm^2^ in control human LMS (p<0.001) (Fig. 5B). Remarkably, this enhancement of cardiac function occurred while maintaining their inherent passive tension (Fig 5B) and contractility kinetics (Fig. 5C), and without triggering ectopic activity (data not shown).

**Figure 5.**
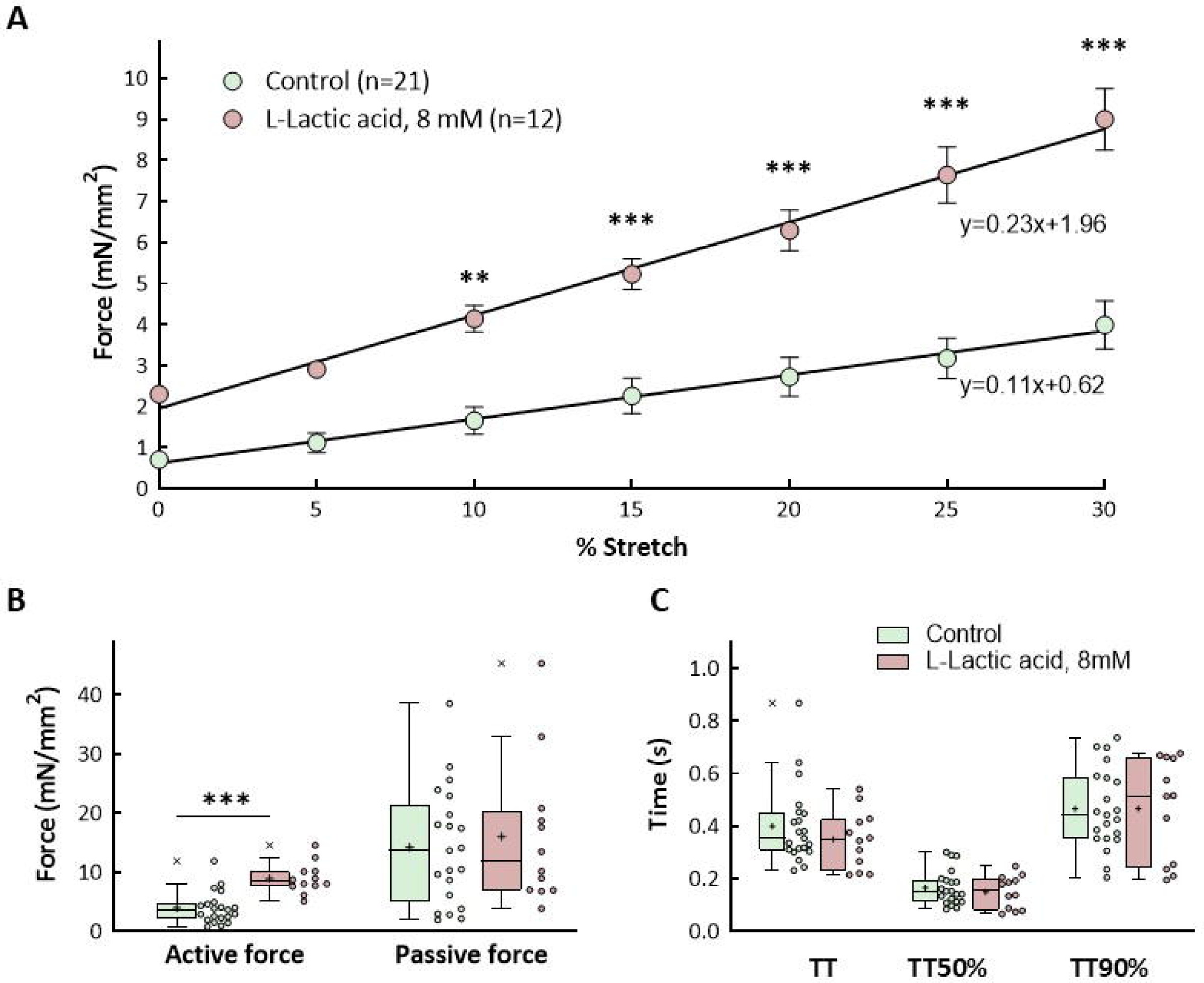
Effect of L-lactic acid on contractility of LMS from human failing hearts. (A) Contraction force (active Frank-Starling relationship) of LMS from human failing hearts under control conditions and after exposure to 8 mmol/L L-lactic acid for 48 hours. ** (p<0.01) and *** (p<0.001) indicate significant differences between both groups (two-way ANOVA followed by *post hoc* Bonferroni test, n=21 and 12 for control and 8 mM lactic acid-treated LMS, respectively). (B) Active and passive force at maximum stretch (i.e., 30%) *** (p<0.001) indicates significant differences between both groups (Student’s t test). (C) Kinetics characteristics of LMS from human failing hearts at maximum contraction, including the time required to reach the peak amplitude of force (TTP, time to peak), and the time to decay from maximum force to 50% (TT50%, time to 50% decay) and to 90% (TT90%, time to 90% decay). Data for B and C are shown as box plot depicting median (horizontal line), mean (+), individual values (color symbols) and outlayers (x).

The effects of L-lactic acid were also characterized in human LMS prepared from two additional donor, healthy, hearts, whose organs where discarded for transplantation. The employment of human LMS from healthy hearts was extremely limited due to its low availability, resulting in small number of samples and reduced statistical power compared to failing human LMS. Nonetheless, L-lactic acid triggered similar effects in healthy adult myocardium as those observed in LMS from failing hearts. L-lactic acid-treated LMS from healthy hearts exhibited enhanced contractility compared to untreated healthy LMS (Supplementary Figure 5).

#### Gene expression

Pooled analysis of gene expression for both LMS from human failing hearts and from two additional donor, healthy, hearts, showed that chronic exposure to exogenous L-lactic acid induced marked changes in gene expression. Myocardial levels of SLC16A3, which encodes for monocarboxylate transporter 4 (MCT4, a high affinity transporter capable of exporting lactate in high-lactate microenvironments ^27^), were significantly enhanced in human LMS exposed to L-lactic acid compared to those found in LMS from human failing hearts under control conditions (Fig. 6). Further, significant upregulations of CACNA1C (encoding for a voltage-dependent calcium channel), CDC25C (involved in cell cycle progression), and the stemness-related gene P53 and the transcription factor OCT4, together with myosin light chain 7 (MYL7) were also observed (Fig. 6). Similar trends were noted for other genes coding for ion channels, proteins involved in cell cycle or acting as transcription factors, but these differences did not reach statistical significance. Separate analysis for samples from LMS from human failing hearts and from LMS from healthy hearts is shown in Supplementary figures 6 and 7. Both types of samples showed similar changes to those previously described.

**Figure 6.**
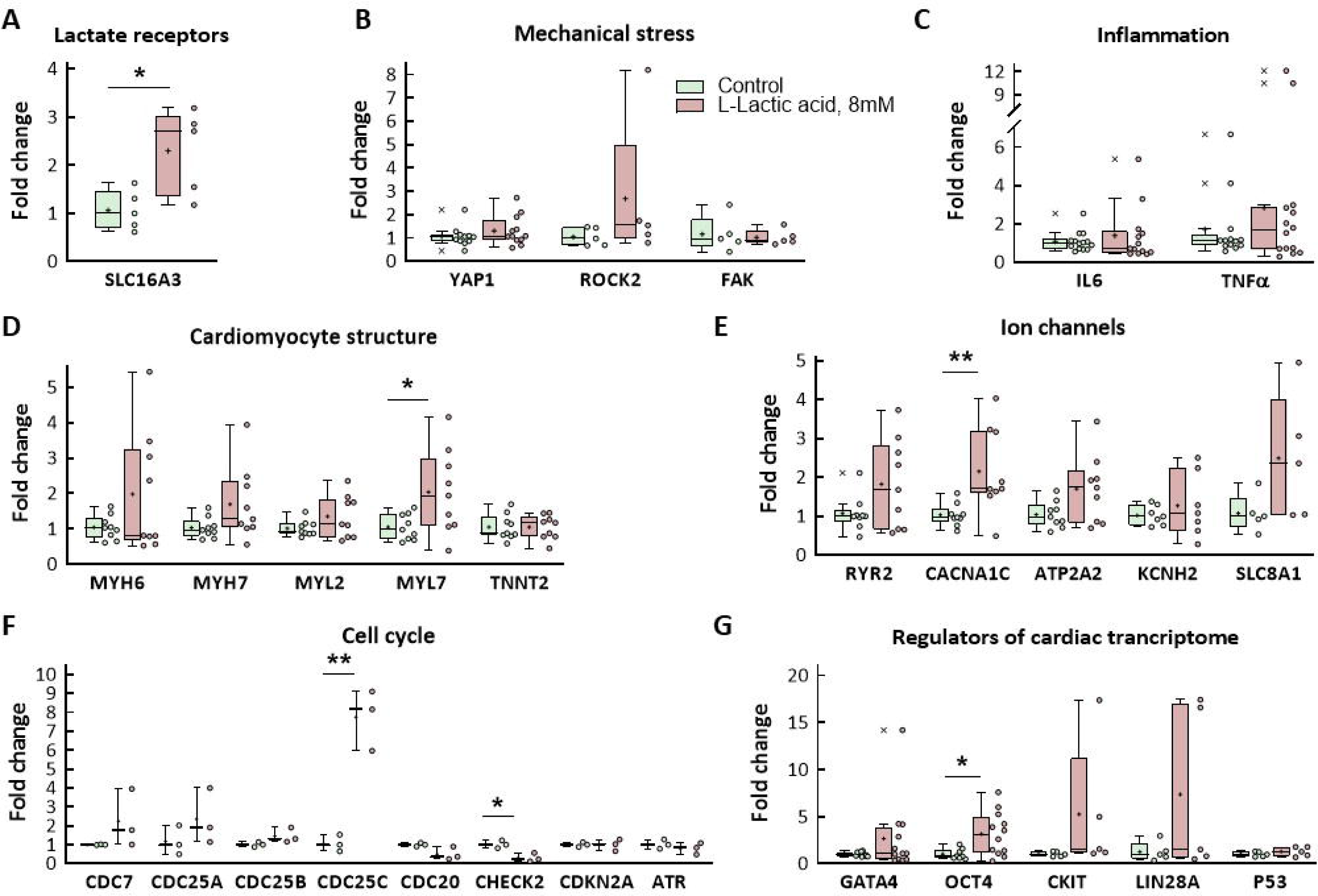
Pooled analysis of gene expression of LMS from both human failing hearts and from two additional donor, healthy, hearts, exposed to control conditions or to incubation with 8 mmol/L L-Lactic acid for 48 hours. (A) Gene encoding for the lactate transporter MCT4. (B) Genes activated by mechanical stress. (C) Inflammatory genes. (D) Cardiomyocyte structural genes. (E) Genes involved in calcium handling. (F) Cell cycle genes. (G) Genes encoding for transcription factors and progenitor genes. * (p<0.05) and ** (p<0.01) indicate significant differences between both groups (Student’s t test) (5 ≤ n ≤ 14). Data are shown as box plot depicting median (horizontal line), mean (+), individual values (color symbols) and outlayers (x).

#### Immunofluorescence analysis of lactate receptors and transporters and angiogenic markers

As depicted in figure 7, expression of MCT4 and of the lactate receptor, GPCR81 (or hydroxycarboxylic acid receptor 1) were significantly enhanced in LMS samples from human failing hearts chronically treated with L-lactic acid. Furthermore, there was a significant upregulation of several proangiogenic markers, including CD31 and the Von Willebrand factor (Fig. 7).

**Figure 7.**
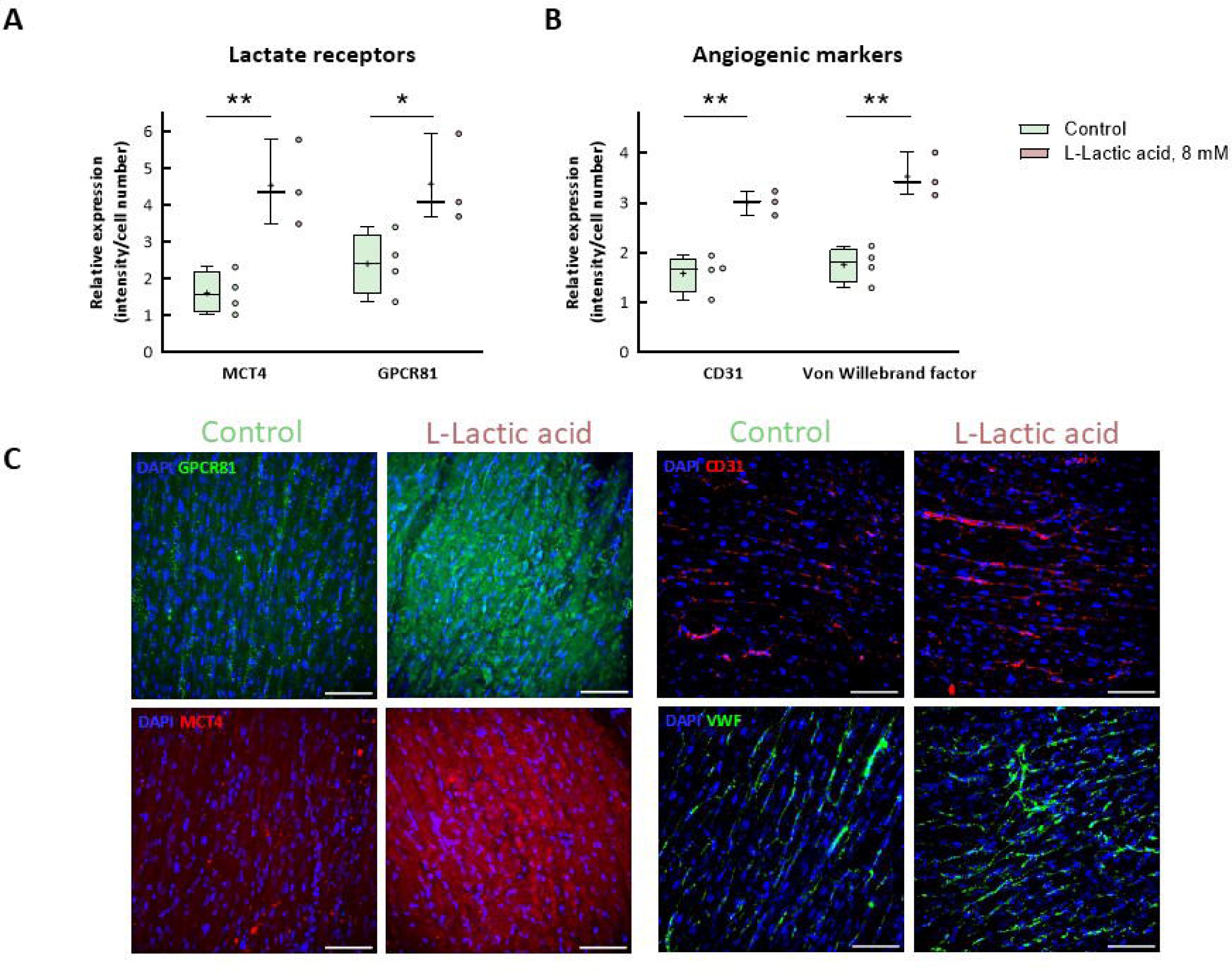
Protein expression of LMS from human failing hearts exposed to control conditions or to incubation with 8 mmol/L L-Lactic acid for 48 hours. Relative expression of (A) lactate receptors and (B) angiogenic markers. * (p<0.05, Student’s t test) indicates significant differences between both groups. Data are shown as box plot depicting median (horizontal line), mean (+), and individual values (color symbols). (C) Representative images of human LMS cultured with control or L-lactic acid-containing medium. Scale bars: 100 μm (n = 4).

## DISCUSSION

This study demonstrates that exogenous L-lactic acid has acute protective effects against reperfusion injury in isolated mice hearts, as denoted by reduced infarct size and LDH release and improved functional recovery at the end of reperfusion. Whereas a significant proportion of these effects can be attributed to the acidotic environment it produces, still an important part can be ascribed to its import through the MCT1 transporter, as demonstrated by attenuation of protection by simultaneous treatment with the MCT-1 inhibitor AR-C 141990. Further, chronic exposure to L-lactic acid enhanced contractility of LMS from human failing hearts without affecting their inherent passive tension and contractility kinetics. These protective actions of L-lactic acid were associated, in the short-term, with acute alterations in the metabolic profile of treated hearts, likely suggesting enhanced substrate availability, and in the long-term, with chronic changes in gene expression, including genes involved in cell cycle progression.

Lactate, a common glycolytic metabolite, has emerged as an important molecule with protective capabilities after brain injuries such as traumatic brain injury ^8,9^ or cerebral ischemia ^10–12^. In these situations, lactate can serve as an alternative energy source, and this occurs not only in the brain, but also in the heart ^9,28–30^. Consequently, here we aimed to explore the potential protective effects of L-lactic acid in adult hearts. Our present data demonstrate that administration of L-lactic acid during initial reperfusion to isolated mice adult hearts results in a concentration-dependent decrease in infarct size, achieving significance at 20 mmol/L. This effect was associated with significant reductions in LDH release and hypercontracture, as well as improved functional recovery at the end of reperfusion. These findings are in contrast with those obtained by others in an in-situ rat model of transient coronary occlusion. In that study, authors were not able to demonstrate any benefit from local, intramyocardial, administration of lactic acid at the onset of reperfusion, despite it induced a reduction in pH in the right atrium and alterations in signaling molecules ^16,17^. These discrepancies might be due to differences in the route and timing of administration.

Addition of 20 mmol/L of L-Lactic acid to the Krebs buffer resulted in a reduction of the buffer’s pH to values near 7.0. It is well known that acidosis during initial reperfusion induces a strong cardioprotective effect ^25,26,31^. Further, a delay in pH recovery from the acidosis that occurs during ischemia may underlie the cardioprotective effects of ischemic postconditioning ^32^. Consequently, it is plausible that a significant part of the observed cardioprotective effect was due to the acidosis induced by L-lactic acid administration. To assess the relative contribution of acidosis to the observed protection, we perfused additional hearts with an acidified Krebs buffer (pH 7.0) during the first 15 min of reperfusion. As expected, treatment with the acidified buffer reduced infarct size compared to control hearts. However, the magnitude of protection obtained with acid buffer was significantly lower than that achieved with L-lactic acid, with the latter exerting a stronger protection than acidosis alone. This result points to the possibility that additional mechanisms are involved in the protective effect of L-lactic acid. Indeed, it has been shown that lactate may be imported into the cytoplasm of cells ^7^, where it may be used as an alternative fuel ^9^, or even exert paracrine effects in neighboring cells or even in distant tissues ^33^.

To assess the potential contribution of transporter-dependent mechanisms in the effect of L-lactic acid, we treated some hearts simultaneously with L-lactic acid and AR-C 141990 hydrochloride, which inhibits the main lactate import system, MCT1 ^7^. Blockade of MCT1 in our experimental setup partially attenuated the protective effect of L-lactic acid, reducing it to a level similar to that found with the acidified Krebs buffer. These data suggest that besides acidosis, L-lactic acid exerts additional protective effects through its uptake by MCT1. In this context, it is worth noting that MCT-1 is a proton-coupled monocarboxylate transporter, so the decrease in pH caused by exogenous lactate promotes its transport inside the cell through the cell membrane.

Once imported into the cytoplasm, lactic acid may serve as a substrate for oxidative phosphorylation and as a major precursor for gluconeogenesis ^6^. The heart is capable of metabolizing various substrates for energy production. The primary fuel of adult cardiomyocytes are fatty acids, but they can easily switch to use carbohydrates, including lactate, and ketone bodies ^2^. Substrate use is influenced by factors as substrate availability, abundance of specific substrate transporters at the cell membrane, or the relative activity of enzymes within metabolic pathways ^2^. Importantly, lactate and fatty acids compete as substrates for cardiac energy metabolism. Particularly, elevated plasma concentrations of lactate are known to reduce fatty acid oxidation ^2,34^. Our present data suggest that L-lactic acid administration triggers a similar metabolic shift in the isolated mouse hearts. Indeed, our ^1^H NMR spectroscopy and discriminant analysis reveal that all four treatment groups have different metabolic profiles, and the pathway and enrichment analysis identified almost a 3-fold enrichment in pyruvate metabolism, glycolysis/gluconeogenesis, and fatty acid biosynthesis, among other pathways altered in L-lactic-treated hearts compared to controls. Remarkably, a metabolic shift towards glycolysis has been previously demonstrated to exert protective actions against cardiac ischemia-reperfusion injury ^35,36^. On the other hand, treatment with acidified buffer resulted in various pathways being altered compared with controls, although with lower significance values.

Further supporting that L-lactic acid may exert protective effects independently of acidosis, recent results have demonstrated that monocytes undergo metabolic reprogramming in the early stage of myocardial infarction and that dysregulated glycolysis and MCT1-mediated lactate transport promote histone lactylation, triggering early activation of the reparative transcriptional response in monocytes. The establishment of the immune homeostasis has been correlated with cardiac repair following myocardial infarction ^37^.

Next, we evaluated the chronic effects of lactate in LMS from adult cardiac tissue. LMS were mainly prepared from the left ventricle of human explanted hearts with distinct cardiomyopathies and exposed to lactate for 48 h. Despite anticipated variability due to inter-individual differences in genetic backgrounds, physiological characteristics, and cardiac pathologies, we were able to show that exposure to L-lactic acid enhanced maximum active force by more than 2-fold compared to control human LMS, thus demonstrating a protective action under these conditions.

Furthermore, chronic exposure to exogenous L-lactic acid induced marked changes in gene expression. Notably, lactic acid-treated human LMS showed a significant upregulation of CACNA1C, encoding for voltage-dependent calcium channels, with similar non-significant trends for other genes encoding for contractile (i.e., MYH6, MYH7, MYL2, MYL7) and Ca^2+^ and K^+^ handling proteins (i.e., RYR2, ATP2A2, KCNH2, SLC8A1). Similarly, others have previously demonstrated an increase in proteins involved in cytoskeletal organization, including actin, myosin, filatin-A or endoplasmin, as well as in phalloidin immunofluoresence, in AC16 cardiomyocytes after 72 h of exposure to lactate^38^.

Significant upregulations were also observed for the stemness-related gene P53, along with the transcription factor OCT4 and with genes involved in cell cycle progression (CDC25C). Similar trends were observed with other genes involved in these processes, as GATA4, CDC7, or CDC25A, although changes did not reach significance due to the high variability related to different donors. GATA4, a critical regulator of cardiac development, differentiation, and cardiomyocyte proliferation ^39,40^, has been shown to induce cardiac differentiation of cardiomyocytes in stem cell-like explants ^41^. Similarly, OCT4, a key factor in the maintenance of an undifferentiated cell state, has been described to have a direct effect on partial cardiomyocyte reprogramming and to enhance the cardiac differentiation potency of progenitor cells ^42,43^. These data may indicate that L-lactic acid induces a shift towards a stem-like phenotype in treated human LMS, similar to the previously described effects in immature cardiac myocytes ^22^. Importantly, similar changes in gene expression were also observed between human LMS from failing hearts and from healthy adult myocardium.

Strikingly, angiogenic proteins, including CD31 or von Willebrand factor, were significantly overexpressed in lactic acid-treated human LMS, supporting the previously described angiogenic potential of lactate ^44^. Additionally, exposure to exogenous lactic acid also increased the expression of lactate receptors at both mRNA and protein levels.

In conclusion, our data support a cardioprotective role for L-lactic acid both in the short-term, following ischemia-reperfusion injury where it reduces infarct size, and in the long-term, in human LMS where it enhanced contractility. While a significant proportion of the acute protective effects can be attributed to maintenance of acidosis early during reperfusion, a role for lactate uptake through the MCT1 transporter is also evident. In the short-term, the protective actions of L-lactic acid were associated with a metabolic shift towards pyruvate metabolism and glycolysis, whereas in the long-term it induced changes in gene expression, including some involved in cell cycle progression. The last findings support the hypothesis of a rejuvenating epigenetic environment for cardiomyocytes, suggesting that lactate could be partially reprogramming cardiomyocytes, activating fetal-like features that would promote injured cardiac regeneration. The favorable effects of L-lactic acid in mature cardiac tissue also endorse its potential use as a scaffold in tissue engineering therapies.

## STUDY LIMITATIONS

A key limitation of this study is the restricted availability of human myocardial tissue samples. Only two heart donors were included, both of whom had died from non-vascular diseases. The employment of human LMS from healthy hearts was extremely limited due to its low availability, resulting in a small number of samples and reduced statistical power compared to studies using failing human LMS. Nonetheless, L-lactic acid triggered similar effects in healthy adult myocardium as those observed in LMS from failing hearts. While this study suggests that the cardioprotective effects of L-lactic acid are consistent across different myocardial conditions, the small sample size of the controls limits the generalizability of these findings. Further studies using a larger and more diverse donor pool are needed to confirm the observed effects and clarify the mechanisms involved in different pathological contexts.

## ACKNOWLEDGEMENTS

This work was supported by the spanish National Research Agency (AEI) (grant BIOCARDIO ref. RTI2018-096320-B-C21) and is part of the I+D+i projects PDC2022-133755-I00 funded by MCIN/AEI/ 10.13039/501100011033 and by “ERDF A way of making Europe” and European Union Next Generation EU. It was also supported by the spanish Instituto de Salud Carlos III (grants PI23/00260 and CIBERCV, from the Spanish Ministry of Science, Innovation and Universities, and being co-financed by the European Regional Development Fund (ERDF-FEDER, a way to build Europe)). Authors are thanked to the Agència de Gestió d’Ajuts Universitaris i de Recerca (2021 SGR 00387) for financial support and from the CERCA program by the Generalitat de Catalunya, and the “Centro de Excelencia Severo Ochoa” (Grant CEX2023-001282-S, funded by MICIU/AEI/ 10.13039/501100011033). Support for the research of E. E. was also received through the prize “ICREA Academia” for excellence in research. This project also received the support of a fellowship from “la Caixa” Foundation (ID 100010434) with fellowship code LCF/BQ/DR19/11740025 and from an EMBO Scientific Exchange Grant (ID 9684) for M. M. LMS work at Imperial College was supported by NC3Rs project grant NC/T001488/1 and BHF studentship to Laura Nicastro (FS/19/57/34894).

## Notes

### Competing Interest Statement

The authors have declared no competing interest.

## REFERENCES

1. Ibanez B, James S, Agewall S, Antunes MJ, Bucciarelli-Ducci C, Bueno H, Caforio ALP, Crea F, Goudevenos JA, Halvorsen S, Hindricks G, Kastrati A, Lenzen MJ, Prescott E, Roffi M, Valgimigli M, Varenhorst C, Vranckx P, Widimský P, Baumbach A, Bugiardini R, Coman IM, Delgado V, Fitzsimons D, Gaemperli O, Gershlick AH, Gielen S, Harjola VP, Katus HA, Knuuti J, Kolh P, Leclercq C, Lip GYH, Morais J, Neskovic AN, Neumann FJ, Niessner A, Piepoli MF, Richter DJ, Shlyakhto E, Simpson IA, Steg PG, Terkelsen CJ, Thygesen K, Windecker S, Zamorano JL, Zeymer U, Chettibi M, Hayrapetyan HG, Metzler B, Ibrahimov F, Sujayeva V, Beauloye C, Dizdarevic-Hudic L, Karamfiloff K, Skoric B, Antoniades L, Tousek P, Shaheen SM, Marandi T, Niemel€a M, Kedev S, Gilard M, Aladashvili A, Elsaesser A, Kanakakis IG, Merkely B, Gudnason T, Iakobishvili Z, Bolognese L, Berkinbayev S, Bajraktari G, Beishenkulov M, Zake I, Lamin H Ben, Gustiene O, Pereira B, Xuereb RG, Ztot S, Juliebø V, Legutko J, Timoteo AT, Tatu-Chiţoiu G, Yakovlev A, Bertelli L, Nedeljkovic M, Studencan M, Bunc M, Castro AMG de, Petursson P, Jeger R, Mourali MS, Yildirir A, Parkhomenko A, Gale CP. 2017 ESC Guidelines for the management of acute myocardial infarction in patients presenting with ST-segment elevation: The Task Force for the management of acute myocardial infarction in patients presenting with ST-segment elevation of the European Society of Cardiology (ESC). Eur Heart J 2018;39:119–177.

2. Zuurbier CJ, Bertrand L, Beauloye CR, Andreadou I, Ruiz-Meana M, Jespersen NR, Kula-Alwar D, Prag HA, Eric Botker H, Dambrova M, Montessuit C, Kaambre T, Liepinsh E, Brookes PS, Krieg T. Cardiac metabolism as a driver and therapeutic target of myocardial infarction. J Cell Mol Med 2020;24:5937–5954.

3. Valls-Lacalle L, Barba I, Miró-Casas E, Alburquerque-Béjar JJ, Ruiz-Meana M, Fuertes-Agudo M, Rodríguez-Sinovas A, García-Dorado D. Succinate dehydrogenase inhibition with malonate during reperfusion reduces infarct size by preventing mitochondrial permeability transition. Cardiovasc Res 2016;109:374–384.

4. Valls-Lacalle L, Barba I, Miró-Casas E, Ruiz-Meana M, Rodríguez-Sinovas A, García-Dorado D. Selective Inhibition of Succinate Dehydrogenase in Reperfused Myocardium with Intracoronary Malonate Reduces Infarct Size. Sci Rep 2018;8.

5. Prag HA, Aksentijevic D, Dannhorn A, Giles A V., Mulvey JF, Sauchanka O, Du L, Bates G, Reinhold J, Kula-Alwar D, Xu Z, Pellerin L, Goodwin RJA, Murphy MP, Krieg T. Ischemia-Selective Cardioprotection by Malonate for Ischemia/Reperfusion Injury. Circ Res 2022;131:528–541.

6. Sun S, Li H, Chen J, Qian Q. Lactic Acid: No Longer an Inert and End-Product of Glycolysis. Physiology (Bethesda) 2017;32:453–463.

7. Li X, Yang Y, Zhang B, Lin X, Fu X, An Y, Zou Y, Wang JX, Wang Z, Yu T. Lactate metabolism in human health and disease. Signal Transduct Target Ther 2022;7.

8. Rice AC, Zsoldos R, Chen T, Wilson MS, Alessandri B, Hamm RJ, Ross Bullock M. Lactate administration attenuates cognitive deficits following traumatic brain injury. Brain Res 2002;928:156–159.

9. Bouzat P, Sala N, Suys T, Zerlauth JB, Marques-Vidal P, Feihl F, Bloch J, Messerer M, Levivier M, Meuli R, Magistretti PJ, Oddo M. Cerebral metabolic effects of exogenous lactate supplementation on the injured human brain. Intensive Care Med 2014;40:412–421.

10. Horn T, Klein J. Neuroprotective effects of lactate in brain ischemia: dependence on anesthetic drugs. Neurochem Int 2013;62:251–257.

11. Berthet C, Lei H, Thevenet J, Gruetter R, Magistretti PJ, Hirt L. Neuroprotective role of lactate after cerebral ischemia. J Cereb Blood Flow Metab 2009;29:1780–1789.

12. Babenko VA, Varlamova EG, Saidova AA, Turovsky EA, Plotnikov EY. Lactate protects neurons and astrocytes against ischemic injury by modulating Ca2+ homeostasis and inflammatory response. FEBS J 2024.

13. Park IH, Cho HK, Oh JH, Chun WJ, Park YH, Lee M, Kim MS, Choi KH, Kim J, Song Y Bin, Hahn JY, Choi SH, Lee SC, Gwon HC, Choe YH, Jang WJ. Clinical Significance of Serum Lactate in Acute Myocardial Infarction: A Cardiac Magnetic Resonance Imaging Study. J Clin Med 2021;10.

14. Montoya JJ, Fernández N, Monge L, Diéguez G, Villalón ÁLG. Nitric oxide-mediated relaxation to lactate of coronary circulation in the isolated perfused rat heart. J Cardiovasc Pharmacol 2011;58:392–398.

15. Groussard C, Morel I, Chevanne M, Monnier M, Cillard J, Delamarche A. Free radical scavenging and antioxidant effects of lactate ion: an in vitro study. J Appl Physiol (1985) 2000;89:169–175.

16. Zhang G, Sun Y, Wang Y, Li X, Li T, Su S, Xu L, Shen H. Local administration of lactic acid and a low dose of the free radical scavenger, edaravone, alleviates myocardial reperfusion injury in rats. J Cardiovasc Pharmacol 2013;62:369–378.

17. Zhang G, Gao S, Li X, Zhang L, Tan H, Xu L, Chen Y, Geng Y, Lin Y, Aertker B, Sun Y. Pharmacological postconditioning with lactic acid and hydrogen rich saline alleviates myocardial reperfusion injury in rats. Sci Rep 2015;5.

18. Koyama T. Postconditioning with Lactate-Enriched Blood for Reducing Lethal Reperfusion Injury in Humans. J Cardiovasc Transl Res 2023;16:793–802.

19. Koyama T, Munakata M, Akima T, Kageyama T, Shibata M, Moritani K, Kanki H, Ishikawa S, Mitamura H. Impact of postconditioning with lactate-enriched blood on in-hospital outcomes of patients with ST-segment elevation myocardial infarction. Int J Cardiol 2016;220:146–148.

20. Castaño O, Pérez-Amodio S, Navarro-Requena C, Mateos-Timoneda MÁ, Engel E. Instructive microenvironments in skin wound healing: Biomaterials as signal releasing platforms. Adv Drug Deliv Rev 2018;129:95–117.

21. Álvarez Z, Castaño O, Castells AA, Mateos-Timoneda MA, Planell JA, Engel E, Alcántara S. Neurogenesis and vascularization of the damaged brain using a lactate-releasing biomimetic scaffold. Biomaterials 2014;35:4769–4781.

22. Ordoño J, Pérez-Amodio S, Ball K, Aguirre A, Engel E. The generation of a lactate-rich environment stimulates cell cycle progression and modulates gene expression on neonatal and hiPSC-derived cardiomyocytes. Biomaterials Advances 2022;139:213035.

23. Watson SA, Scigliano M, Bardi I, Ascione R, Terracciano CM, Perbellini F. Preparation of viable adult ventricular myocardial slices from large and small mammals. Nat Protoc 2017;12:2623–2639.

24. Watson SA, Duff J, Bardi I, Zabielska M, Atanur SS, Jabbour RJ, Simon A, Tomas A, Smolenski RT, Harding SE, Perbellini F, Terracciano CM. Biomimetic electromechanical stimulation to maintain adult myocardial slices in vitro. Nat Commun 2019;10.

25. Rodriguez-Sinovas A, Cabestrero A, Garcia del B, Inserte J, Garcia A, Garcia-Dorado D. Intracoronary acid infusion as an alternative to ischemic postconditioning in pigs. Basic Res Cardiol 2009;104:761–771.

26. Inserte J, Barba I, Hernando V, Abellán A, Ruiz-Meana M, Rodríguez-Sinovas A, Garcia-Dorado D. Effect of acidic reperfusion on prolongation of intracellular acidosis and myocardial salvage. Cardiovasc Res 2008;77:782–790.

27. Contreras-Baeza Y, Sandoval PY, Alarcón R, Galaz A, Cortés-Molina F, Alegriá K, Baeza-Lehnert F, Arce-Molina R, Guequén A, Flores CA, Martín AS, Barros LF. Monocarboxylate transporter 4 (MCT4) is a high affinity transporter capable of exporting lactate in high-lactate microenvironments. J Biol Chem 2019;294:20135–20147.

28. Brooks GA. Cell-cell and intracellular lactate shuttles. J Physiol 2009;587:5591–5600.

29. Brooks GA. The Science and Translation of Lactate Shuttle Theory. Cell Metab 2018;27:757–785.

30. Brooks GA. Role of the Heart in Lactate Shuttling. Front Nutr 2021;8.

31. Lemasters JJ, Bond JM, Chacon E, Harper IS, Kaplan SH, Ohata H, Trollinger DR, Herman B, Cascio WE. The pH paradox in ischemia-reperfusion injury to cardiac myocytes. EXS 1996;76:99–114.

32. Inserte J, Ruiz-Meana M, Rodríguez-Sinovas A, Barba I, Garcia-Dorado D. Contribution of delayed intracellular pH recovery to ischemic postconditioning protection. Antioxid Redox Signal 2011;14:923–939.

33. Pellerin L, Pellegri G, Bittar PG, Charnay Y, Bouras C, Martin JL, Stella N, Magistretti PJ. Evidence supporting the existence of an activity-dependent astrocyte-neuron lactate shuttle. Dev Neurosci 1998;20:291–299.

34. Vusse GJ van der, Groot MJM de. Interrelationship between lactate and cardiac fatty acid metabolism. Mol Cell Biochem 1992;116:11–17.

35. Alburquerque-Béjar JJ, Barba I, Inserte J, Miró-Casas E, Ruiz-Meana M, Poncelas M, Vilardosa Ú, Valls-Lacalle L, Rodríguez-Sinovas A, Garcia-Dorado D. Combination therapy with remote ischaemic conditioning and insulin or exenatide enhances infarct size limitation in pigs. Cardiovasc Res 2015;107:246–254.

36. Beltran C, Pardo R, Bou-Teen D, Ruiz-Meana M, Villena JA, Ferreira-González I, Barba I. Enhancing Glycolysis Protects against Ischemia-Reperfusion Injury by Reducing ROS Production. Metabolites 2020;10.

37. Wang N, Wang W, Wang X, Mang G, Chen J, Yan X, Tong Z, Yang Q, Wang M, Chen L, Sun P, Yang Y, Cui J, Yang M, Zhang Y, Wang D, Wu J, Zhang M, Yu B. Histone Lactylation Boosts Reparative Gene Activation Post-Myocardial Infarction. Circ Res 2022;131:893–908.

38. Luti S, Militello R, Pinto G, Illiano A, Marzocchini R, Santi A, Becatti M, Amoresano A, Gamberi T, Pellegrino A, Modesti A, Modesti PA. Chronic lactate exposure promotes cardiomyocyte cytoskeleton remodelling. Heliyon 2024;10.

39. Rojas A, Kong SW, Agarwal P, Gilliss B, Pu WT, Black BL. GATA4 is a direct transcriptional activator of cyclin D2 and Cdk4 and is required for cardiomyocyte proliferation in anterior heart field-derived myocardium. Mol Cell Biol 2008;28:5420–5431.

40. Afouda BA. Towards Understanding the Gene-Specific Roles of GATA Factors in Heart Development: Does GATA4 Lead the Way? Int J Mol Sci 2022;23.

41. Latinkić B V., Kotecha S, Mohun TJ. Induction of cardiomyocytes by GATA4 in Xenopus ectodermal explants. Development 2003;130:3865–3876.

42. Yannarelli G, Pacienza N, Montanari S, Santa-Cruz D, Viswanathan S, Keating A. OCT4 expression mediates partial cardiomyocyte reprogramming of mesenchymal stromal cells. PLoS One 2017;12.

43. Nagura S, Otaka S, Koike C, Okabe M, Yoshida T, Fathy M, Fukahara K, Yoshimura N, Misaki T, Nikaido T. Effect of exogenous Oct4 overexpression on cardiomyocyte differentiation of human amniotic mesenchymal cells. Cell Reprogram 2013;15:471–480.

44. Zhou J, Liu T, Guo H, Cui H, Li P, Feng D, Hu E, Huang Q, Yang A, Zhou J, Luo J, Tang T, Wang Y. Lactate potentiates angiogenesis and neurogenesis in experimental intracerebral hemorrhage. Exp Mol Med 2018;50.

