## Supplemental for "L-lactic acid induces short- and long-term cardioprotective effects through MCT1 transport, induction of metabolic reprogramming, and gene expression modulation"

**Supplementary methods.**

Animal studies complied with European legislation (Directive 2010/63/EU) on the protection of animals used for scientific purposes, with the Guide for the Care and Use of Laboratory Animals published by the US National Institutes of Health (NIH Publication No. 85-23, revised 1996, updated in 2011), and were approved by the Ethics Commettee of Vall d’Hebron Research Institute (CEEA31.15). The use of human samples adhered to the principles outlined in the Declaration of Helsinki and followed the European Guidelines for Good Clinical Practice. Human tissues from patients with heart failure and from two cardiac transplant donors whose organs were discarded for transplantation were provided by the NIHR Cardiovascular Biomedical Research Unit (Royal Brompton & Harefield NHS Foundation Trust and Imperial College London). The study received approval from the UK institutional ethics committee (NRES ethics number for biobank samples: 09/H0504/104 + 5; Biobank approval number: NP001-06-2015 and MED_CT_17_079) and Imperial College London. Informed consent was obtained from all patients or their families.

**Isolated, Langendorff-perfused, mice hearts.**

Adult male C57BL/6J mice (25-30g, 9-12 weeks) were used throughout the study. Animals were euthanized with an intraperitoneally overdose of sodium pentobarbital (1.5 g/Kg). Immediately, mice underwent a bilateral thoracotomy and the whole hearts were quickly removed. Hearts were, then, retrogradely perfused through the aorta with an oxygenated (95% O_2_: 5% CO_2_) Krebs solution (in mmol/L: NaCl 118, KCl 4.7, MgSO_4_ 1.2, CaCl_2_ 1.8, NaHCO_3_ 25, KH_2_PO_4_ 1.2, and glucose 11, pH 7.4), at 37°C, using a constant flow Langendorff system, as previously described ^1^. Flow was initially adjusted to produce a perfusion pressure of 80-90 mmHg (normoxic conditions; i.e., about 3.5 mL/min). Left ventricular (LV) pressure was monitored with a water-filled latex balloon connected to a pressure transducer, placed in the left ventricle and inflated to obtain a LV end-diastolic pressure (LVEDP) between 6 and 8 mmHg.

*Concentration–response curves to L-lactic acid during normoxia.*

After a 30-minute equilibration period, mouse hearts were treated, under normoxic conditions, with Krebs buffer containing increasing concentrations of L-(+)-Lactic acid (#199257, Sigma-Aldrich, St. Louis, USA), ranging from 1 to 50 mmol/L. Each lactic acid concentration was directly dissolved in freshly prepared Krebs buffer and given for 10 minutes, while maintaining oxygenation throughout the experiment. Osmolarity was maintained constant by addition of D-(+)-sucrose (PanReac AppliChem, Castellar del Vallès, Spain) to match that of the highest lactic acid concentration. The pH of the solutions was adjusted to 7.4. Lactate dehydrogenase (LDH) release was determined at the end of each concentration from samples from the coronary effluent, and infarct size was measured at end of the experiment by TTC staining. Concentration-response curves for left ventricular developed pressure (LVdevP), left ventricular end diastolic pressure (LVEDP), heart rate, and perfusion pressure were fitted to sigmoid curves using the equation y=y0+a/[1+exp(-x-x0)/b], to determine the concentration causing half-maximal effect (EC50) and the maximal effect (Emax). LVdevP was defined as the systolic pressure minus LVEDP.

*Effects of L-lactic acid on myocardial ischemia–reperfusion injury.*

Following a 30-minute stabilisation period, mouse hearts were subjected to 35 minutes of global ischaemia followed by 60 minutes of reperfusion, as described previously ^1^. During ischaemia, hearts were immersed in saline solution to maintain temperature at 37 °C. Hearts were, then, allocated to the different experimental groups. Control hearts (n=7) received the standard Krebs buffer during the entire reperfusion (pH 7.4), whereas L-lactic acid-treated hearts (n=5/group) received Krebs containing L-(+)-Lactic acid at 8 or 20 mmol/L, beginning at the onset of reperfusion and for the its first 15 minutes. Selected concentrations were in the same order of magnitude as the EC50 found in the normoxic experiments.

As pH of L-lactic acid-containing solutions was about 7.0, and to assess the relative contribution of acidosis to the observed effects, additional hearts were perfused with an acidic Krebs buffer (in mmol/L: NaCl 118, KCl 4.7, MgSO_4_ 1.2, CaCl_2_ 1.8, NaHCO_3_ 7.5, KH_2_PO_4_ 1.2, and glucose 11, pH 7.0) during the first 15 min of reperfusion (n=5) and results were compared with those obtained with 20 mmol/L L-lactic acid. In addition, to assess whether the effects of L-lactic acid were dependent on its transport into the cytoplasm, some hearts were simultaneously perfused with Krebs buffer containing 20 mmol/L L-(+)-Lactic acid during the first 15 min of reperfusion while inhibiting its main transport system, the monocarboxylate transporter 1 (MCT1) ) ^2^, with AR-C141990 hydrochloride 1 μmol/L (#5658, Tocris, Bristol, UK) (n=6).

Functional recovery (as a percentage of the baseline LVdevP), heart rate and hypercontracture (the difference between the maximum LVEDP value during the initial minutes of reperfusion and the value at the end of ischaemia) were assessed in all cases.

*LDH release and infarct size measurements.*

Cell death was indirectly quantified by measuring LDH release in samples from the coronary effluent by spectrophotometry, as previously described ^1^. Infarct size was determined in cardiac slices by 2,3,5-triphenyltetrazolium chloride staining at the end of reperfusion ^1^.

*Analysis of myocardial metabolism.*

Cardiac metabolites were analyzed in 24 additional mouse hearts subjected to 35 minutes of global ischemia followed by 5 minutes of reperfusion, under the different treatments previously described (n=6/group). Metabolites from the cardiac ventricles were extracted using the methanol:chloroform method ^1^. Extracts were dissolved in 600 ml of deuterium oxide containing 1 mmol/L of TSP (3-(trimethylsilyl)propionic-2,2,3,3-d4 acid) and ^1^H NMR spectra were acquired on a vertical bore 9.4T magnet interfaced to a Bruker Avance 400 spectrometer ^1^. One-way analysis of variance (ANOVA) and Fisher LSD post-hoc tests and OPLS-discriminant analysis were performed using SIMCA 14.0 (Umetrics, Sartorius, Germany). Metabolites were matched to metabolomics pathways using the Pathway Analysis and Enrichment Analysis features in Metaboanalyst 6.0.

**Human living myocardial slices (LMS).**

Hearts from six patients with heart failure and from two donors who died from non-cardiovascular diseases were isolated, placed in cold cardioplegia solution, and stored in ice. Patient characteristics are detailed in supplementary table 1. Human LMS were quickly prepared as previously described ^3^. In brief, a tissue block of ~1.5 cm^2^ was obtained from the left ventricular wall and mounted, with the epicardial surface facing down, onto an agarose-coated specimen holder, glued to its surface with Histoacryl surgical glue. The specimen holder was then submerged in a vibratome bath containing cold (4 °C), oxygenated Tyrode's slicing solution (in mmol/L: NaCl 140, KCl 9, MgCl_2_ 1, CaCl_2_ 1, HEPES 10, glucose 10, and BDM 30, pH 7.4). A high-precision vibrating microtome (7000 smz-2; Campden Instruments, London, UK) with a ceramic blade was employed to prepare the human LMS using the following setup: vibrating frequency 80 Hz, thickness 300 µm, amplitude 2 mm, and advance speed 0.03 mm/s. The tissue block was sliced from the endocardium, in parallel to the myofibers’ direction.

*LMS culture.*

Square cardiac slices (~10x10mm) with homogeneous fiber alignment, were cut from these sections using a razor blade. Polyethylene terephthalate (Taulman3D T-glase) custom-made, biocompatible, rectangular holders were 3D-printed and glued along the width of the slice, in a direction perpendicular to the fibers. Henceforth, handling of LMS was carried out in a sterile laminar flow cabinet. The slices were, finally, stretched to a physiological load (i.e., sarcomere length of 2.2 µm ^4^) by sterile, custom, stainless-steel stretchers. The percentage of stretch to mimic pre-load conditions in vitro for human LMS from failing hearts was 19.6% ^4^.

After LMS preparation and stretch, slices were immediately cultured *in vitro* in custom sealed culture chambers, at 37ºC, under continuous oxygenation (95% O_2_: 5% CO_2_). Culture medium was continuously recirculated, and consisted of medium 199 (Sigma-Aldrich, St. Louis, USA) supplemented with (in nmol/L) adrenaline 4, noradrenaline 4, dexamethasone 100, and 3,3′,5-Triiodo-L-thyronine 2.15, plus insulin-transferrin-selenium 0.1%, penicillin-streptomycin 2%, and ascorbic acid 20 μg/mL. LMS preparations were electrically stimulated using carbon electrodes at 0.5 Hz (10 ms pulse width, 15 V). LMS were randomly assigned to control or L-lactic acid-treated groups. L-(+)-lactic acid was directly dissolved in the cell culture medium at a concentration of 8 mmol/L. This concentration is in the same order of magnitude of the EC50 found in normoxic mice hearts, but slightly lower than that used in ischemia-reperfusion experiments. The reduced concentration was based on previous optimization studies conducted with human cardiac cells *in vitro*, which revealed cytotoxic effects at concentrations above 8 mmol/L. Additionally, Ordoño *et al.* ^5^ showed that human cardiac cells exhibit greater sensitivity to lactate in comparison to rodent cells.

*LMS contractility.*

The contraction force was measured immediately after cultivation using a force transducer (HSE isometric force transducer F30 type 372, Harvard Apparatus, USA). Cardiac slices were affixed to the force transducer using the 3D-printed holders, and stimulated at 0.5 Hz (10 ms pulse width, 20–30 V). The slices were stretched gradually, in a stepwise manner, until maximum isometric contraction was achieved. Force signals and peak amplitudes were recorded and analyzed using AxoScope and Clampfit software (Molecular Devices, San Jose, USA). Developed forces were normalized to the cross-sectional area and reported as wall stress (mN/mm^2^).

*Gene expression.*

Quantitative real-time reverse transcription-polymerase chain reaction (qRT-PCR) was used to investigate the relative expression levels of various cardiac genes. Following culture with or without L-lactic acid, LMS were washed with PBS, detached from the holders with a razor blade, and submerged in liquid nitrogen. Snap-frozen LMS were stored at -20 °C until mRNA isolation.

Frozen LMS samples were placed in a pre-cooled tube containing a 5 mm stainless-steel bead (Qiagen) and cold TRIzol^TM^ (Invitrogen^TM^) and lysed with a TissueLyser LT (Qiagen) at 40 Hz, for 3–5 minutes. Next, 100 μL of chloroform (Sigma-Aldrich, St. Louis, USA) was added and the tubes were kept at RT for 5 minutes with intermittent shaking. The tubes were, then, centrifuged at 12000 rpm, for 15 minutes at 4 °C, resulting in phase separation. The top aqueous layer, containing the RNA, was carefully transferred to a clean tube, and an equal volume of 70% ethanol was added and mixed thoroughly. Finally, the solution was transferred to a RNeasy spin column and RNA was extracted with a RNeasy® Mini Kit (Qiagen) according to manufacturer’s instructions.

RNA purity and concentration were determined using a NanoDrop Spectrophotometer (ND-1000 Spectrophotometer, NanoDrop®, Thermo Fisher Scientific, Waltham, USA). For each sample, 500 ng of total RNA was reverse transcribed to cDNA using iScript™ cDNA Synthesis Kit (Bio-Rad, Hercules, USA). A no reverse transcription control was used by replacing the iScript Reverse Transcriptase by nuclease-free water. cDNA (12.5 ng) was mixed with iTaq™ Universal SYBR® Green Supermix (Bio-Rad, Hercules, USA) and a primers mix (containing 10 μmol/L of forward and reverse primers) (Supplementary Table 1). Quantitative real-time PCR was carried out in a QuantStudio 6 Pro Real-time PCR system (Applied Biosystems, Waltham, USA). The following run settings were used: 30’’ at 95 °C (polymerase activation), followed by 40 cycles of 15’’ at 95 °C and 60’’ at 60 °C (denaturation-extension), followed by a melt curve analysis (instrument default settings). Relative gene-fold variations were calculated by the 2^-ΔΔCt^ method using GAPDH as the housekeeping gene. Non-treated LMS (without L-lactic acid) were considered as control samples. Three technical replicates per sample were measured and the resulting Ct values averaged.

*Immunofluorescence staining.*

After culture with and without L-lactic acid, LMS were washed with PBS, fixed in 4% formaldehyde solution (Thermo Scientific Scientific, Waltham, USA) for 15 min, and washed again in PBS. Fixed LMS were then permeabilized and non-specific binding blocked in 1% (v/v) Triton X–100 (Sigma-Aldrich, St. Louis, USA), 10% FBS (Gibco, Thermo Fisher Scientific, Waltham, USA), 5% (w/v) BSA (Sigma-Aldrich, St. Louis, USA) and 10% (v/v) horse serum (Gibco, Thermo Fisher Scientific, Waltham, USA) in PBS solution for 3 hours at room temperature. LMS were then washed and incubated overnight at 4ºC with primary antibodies dissolved in PBS- 1% BSA (mouse anti-CD31/PECAM-1, #sc-376764, Santa Cruc Biotechnology, dilution 1:50; rabbit anti-GPCR GPR81, #orb183872, Biorbyt, dilution 1:100; rabbit anti-SLC16A3 (MCT4), #orb6971, Biorbyt, dilution 1:100; mouse anti-Von Willebrand factor, #sc-53466, Santa Cruz Biotechnology, dilution 1:100). Samples were then washed three times with PBS, for 30 minutes, and incubated for 2h (RT) with the corresponding goat secondary anti-mouse or anti-rabbit antibodies marked with Alexa Fluor 488 or 568 (#ab150113, #ab175473, #ab150077 (dilution 1:500) or #ab175471 (dilution 1:1000)). Lastly, LMS were counterstained with 4′,6-diamidino-2-phenylindole (DAPI) (Sigma-Aldrich, St. Louis, USA, #D9542, dilution 1:500) for 15 min, washed three times for 15 min, and stored at 4 °C until imaged. Immunolabelled samples were imaged using a wide-field epifluorescence microscope (Leica Thunder 3D Live Cell, Leica Biosystems Nussloch GmbH, Heidelberger, Germany). Quantitative image analysis was performed using Fiji software. A minimum of 6 random images per sample were acquired and analyzed, and the result averaged.

*Statistical analyses.*

Results are shown as mean ± standard error (SEM). Statistical differences were determined using GraphPad Prism 9.4.0. For two group comparisons, a two-tailed Student’s T-test was used. For multiple comparisons, one-way ANOVA followed by *post hoc* Tukey’s test (infarct size, cumulative LDH release, hypercontracture) or two-way ANOVA followed by *post hoc* Bonferroni test (Frank-Starling relationship curves) were performed. Repeated measures ANOVA (MANOVA) followed by *post hoc* Tukey’s test was used for analysis of one intergroup and one intragroup variable (changes in LVdevP, heart rate, LVEDP, perfusion pressure). Differences were considered significant when p<0.05.

**Supplementary Table 1.** Characteristics of healthy donors and of patients with pathological failing hearts from whom LMS preparation were obtained.

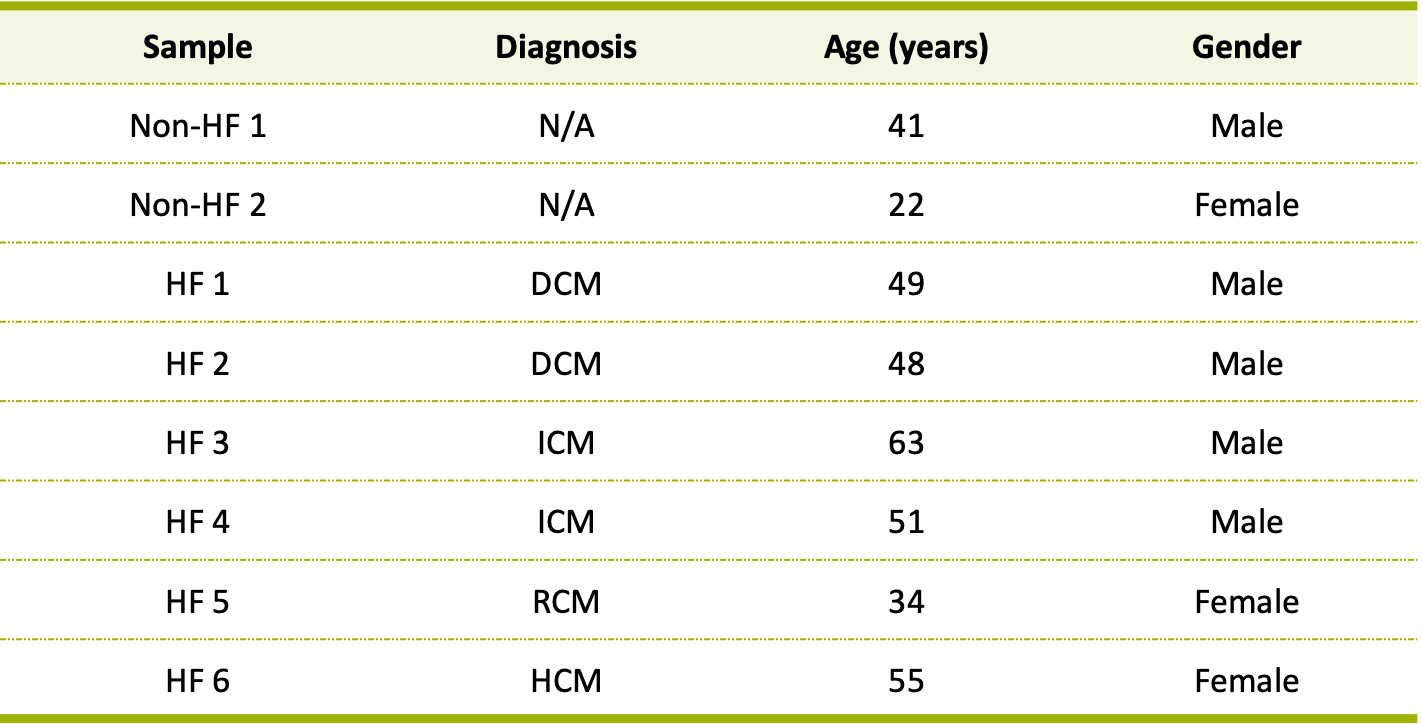

HF – Heart failure; DCM – Dilated cardiomyopathy; ICM – Ischemic cardiomyopathy; RCM – Restrictive cardiomyopathy; HCM – Hypertrophic cardiomyopathy; N/A – Not applicable.

**Supplementary Table 2.** List of primers used for cDNA amplification by RT-qPCR in LMS samples from human hearts.

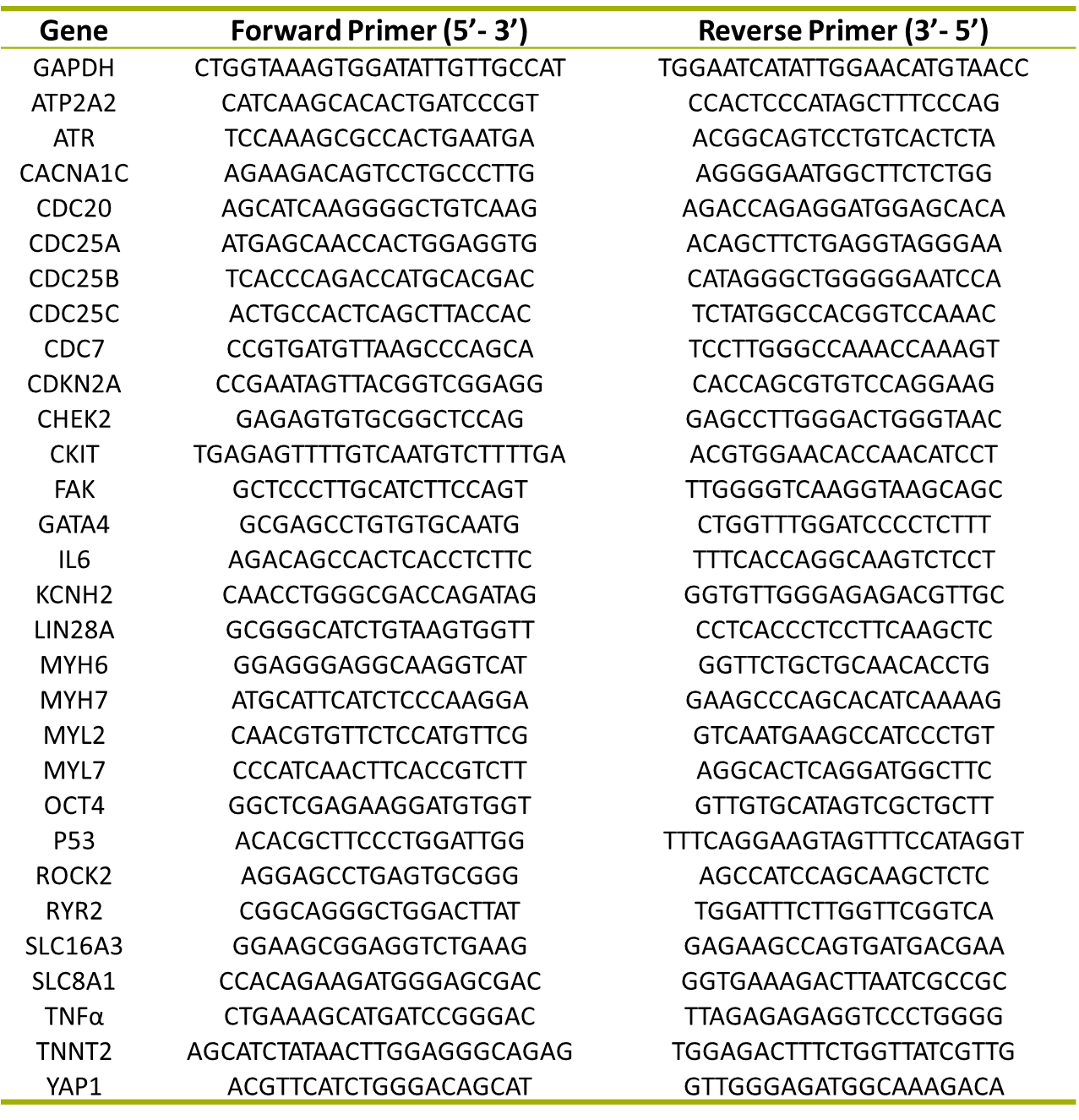

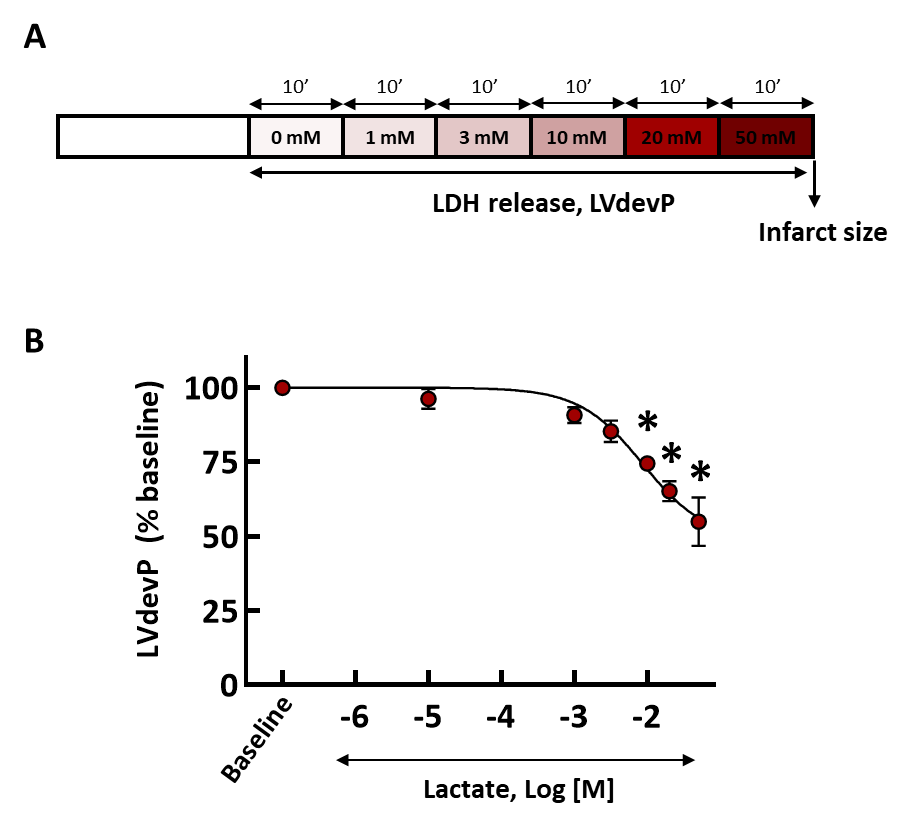

**Supplementary Figure 1.** (A) Experimental design used to assess the effects of different concentrations of L-lactic acid, ranging from 1 to 50 mmol/L in isolated mice hearts. Each concentration was applied for 10 minutes. (B) Concentration-response curve for left ventricular developed pressure (LVdevP) in normoxic mice hearts perfused with increasing concentrations of L-lactic acid (n=4). * (p<0.05) indicates significant differences respect to baseline values (repeated measures ANOVA and Tukey’s post hoc tests).

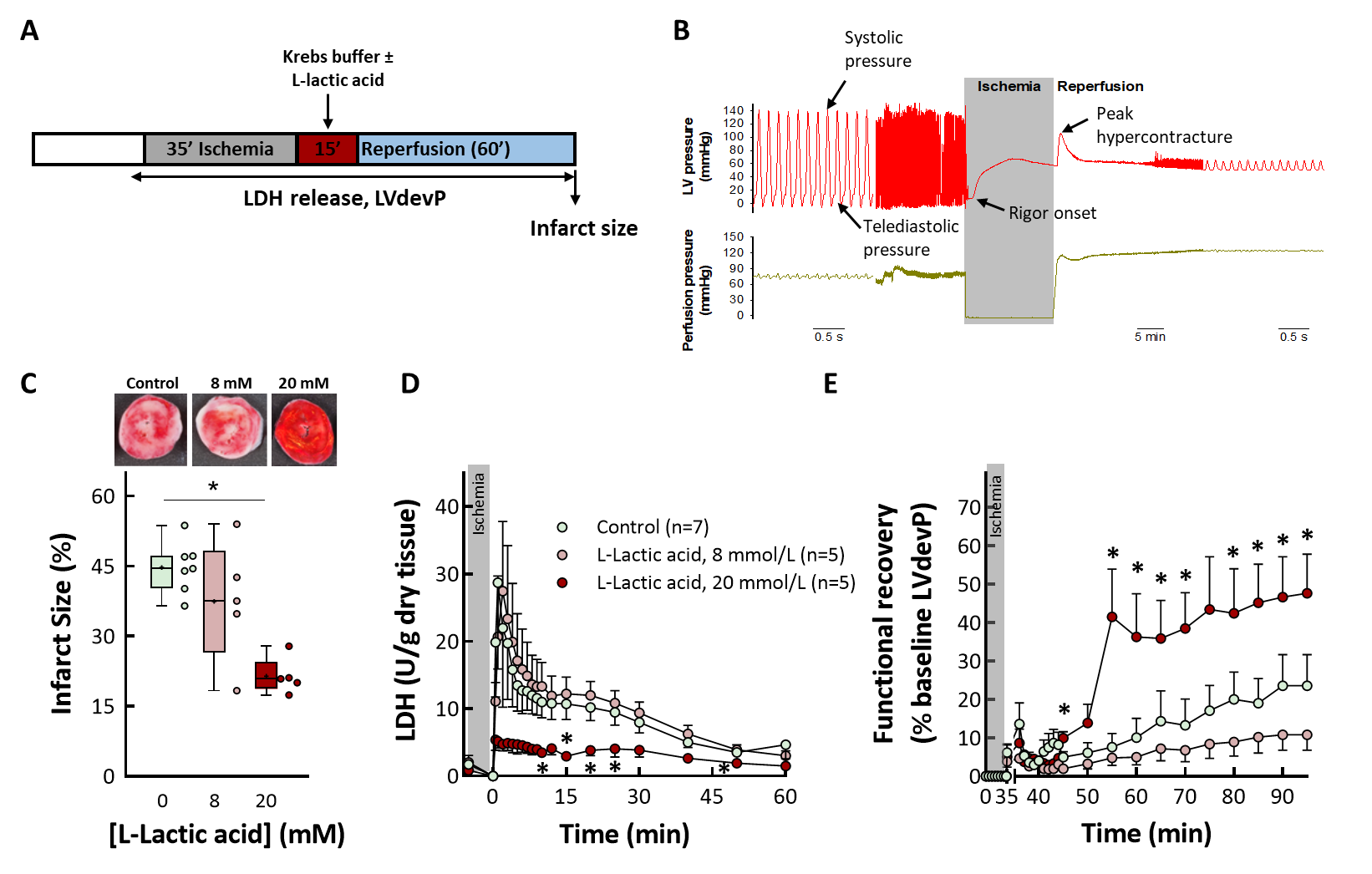

**Supplementary Figure 2.** (A) Experimental design used to test the effects of L-lactic acid, given at the onset of reperfusion, in isolated mice hearts submitted to 35 min global ischemia followed by reperfusion. Experimental design of ischaemia-reperfusion experiments. (B) Representative recording showing changes induced by ischemia and reperfusion on left ventricular pressure and perfusion pressure in a mice heart included in the study. (C) Effects of 8 and 20 mmol/L of L-lactic acid (n=5) on myocardial infarct size as compared with control hearts (n=7). * (p<0.05) indicates significant differences respect to baseline values (ANOVA and Tukey’s post hoc tests). Data are shown as box plot depicting median (horizontal line), mean (+), and individual values (color symbols). (D) Effects on cumulative LDH release. * (p<0.05) indicates significant differences respect to values in the control group (repeated measures ANOVA and Tukey post-hoc tests). (E) Functional recovery in mice hearts submitted to 35 min global ischemia followed by reperfusion and treated or not with L-lactic acid (8 and 10 mmol/L). * (p<0.05) indicates significant differences respect to values in the control group (repeated measures ANOVA and Tukey post-hoc tests).

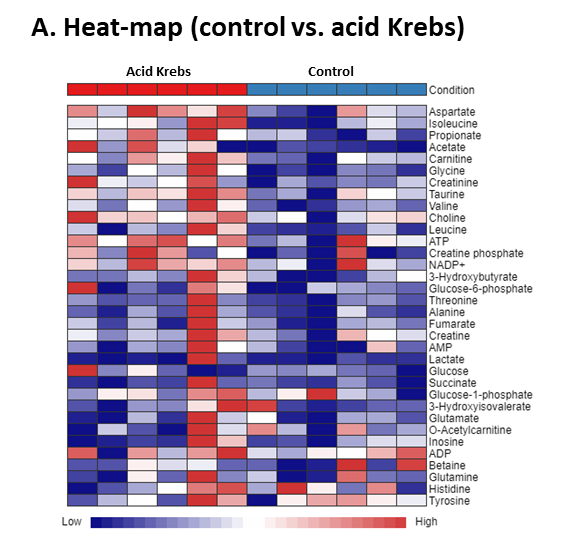

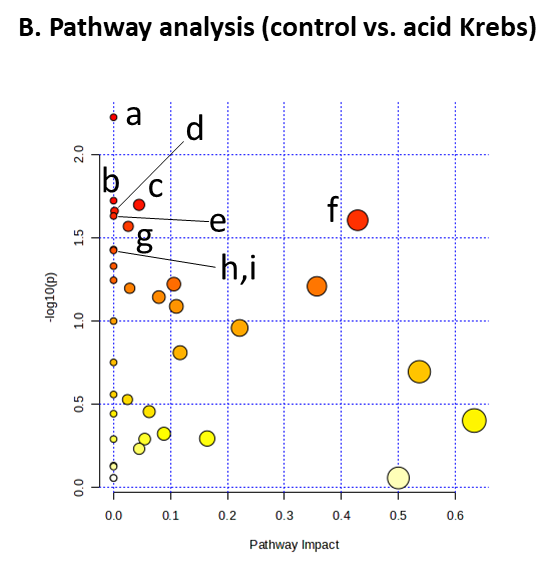

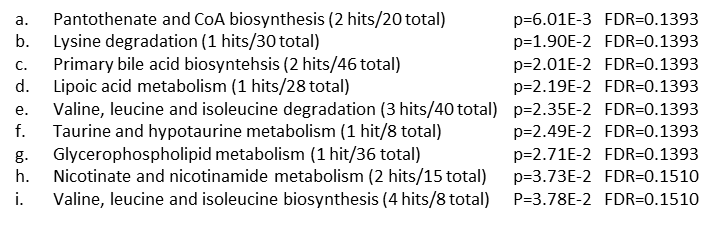

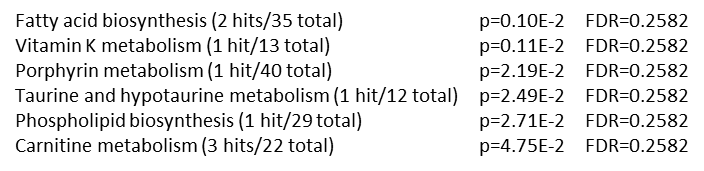

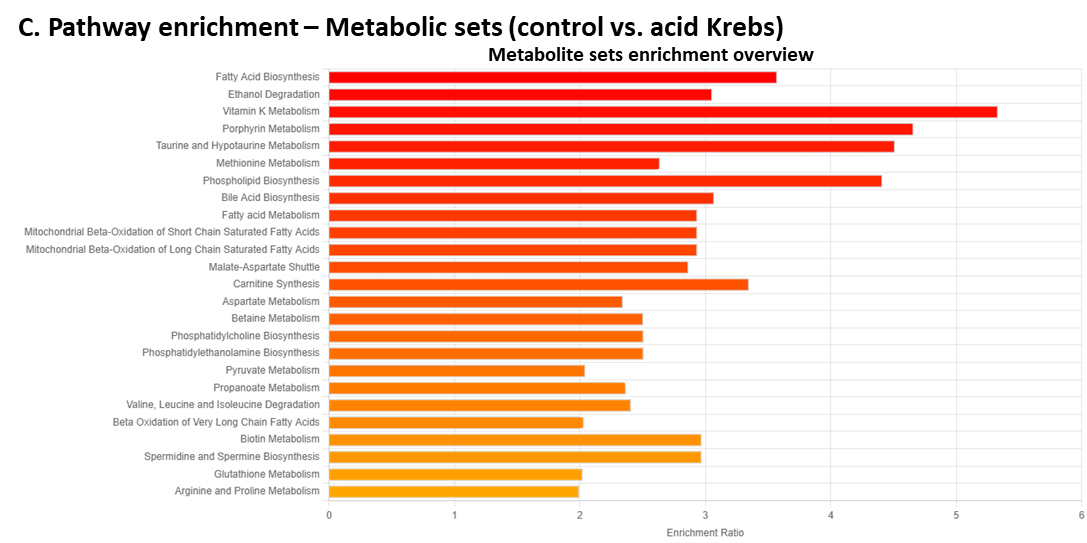

**Supplementary Figure 3.** Non-targeted metabolomic analysis of hearts from mice hearts submitted to 35 min global ischemia followed by reperfusion, and treated, during the first 15 min of reperfusion, with standard Krebs (n=7), or with an acidic Krebs buffer (pH 7.0) (n=5). (A) Heat-map of metabolites between individual cases in control and L-lactic acid-treated hearts. (B) Pathway analysis of analyzed metabolites. (C) Pathway enrichment analysis of analyzed metabolites using metabolic datasets.

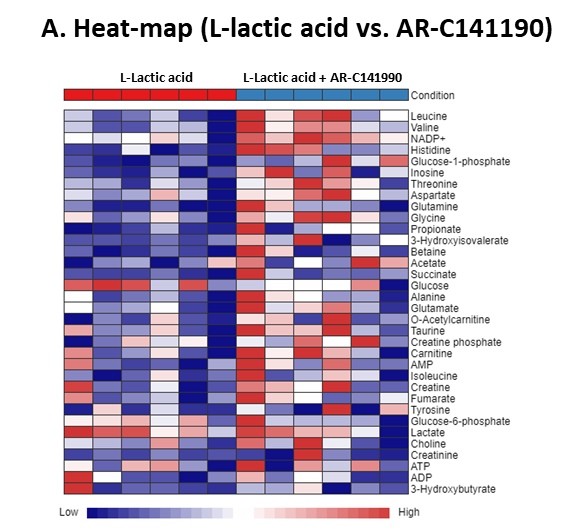

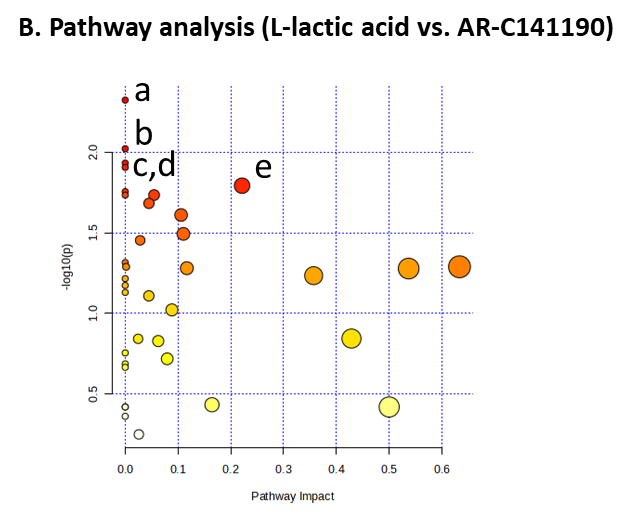

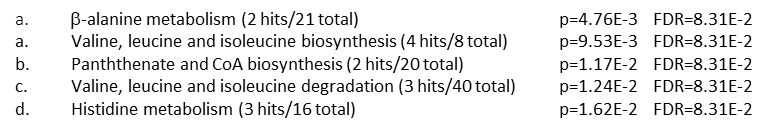

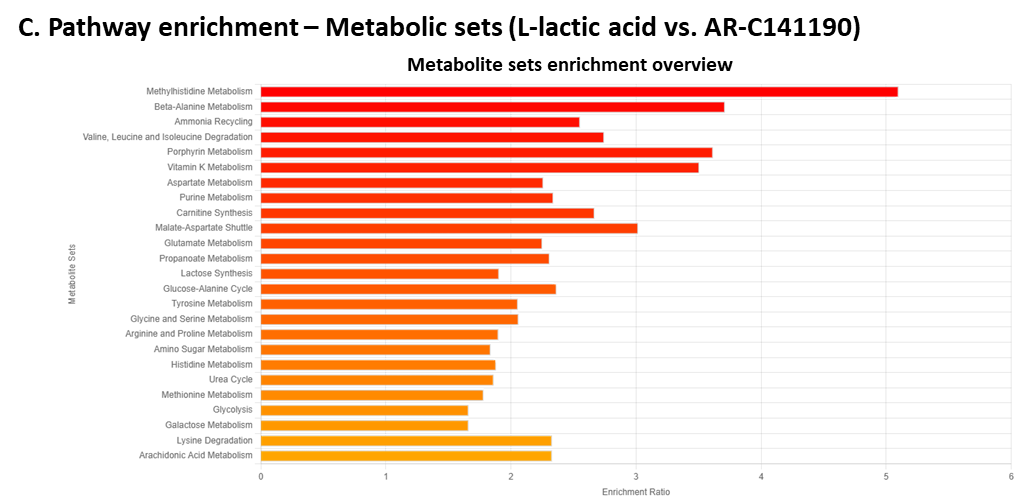

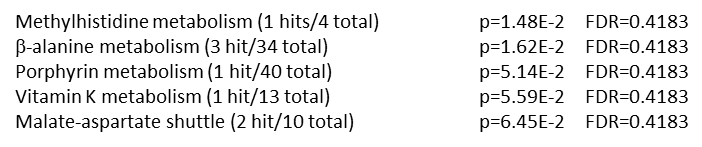

**Supplementary Figure 4.** Non-targeted metabolomic analysis of hearts from mice hearts submitted to 35 min global ischemia followed by reperfusion, and treated, during the first 15 min of reperfusion, with Krebs containing L-lactic acid (n=5) or L-lactic acid supplemented with AR-C141990. (A) Heat-map of metabolites between individual cases in both groups. (B) Pathway analysis of analyzed metabolites. (C) Pathway enrichment analysis of analyzed metabolites using metabolic datasets.

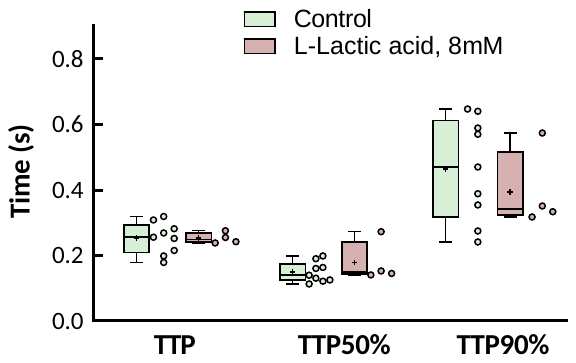

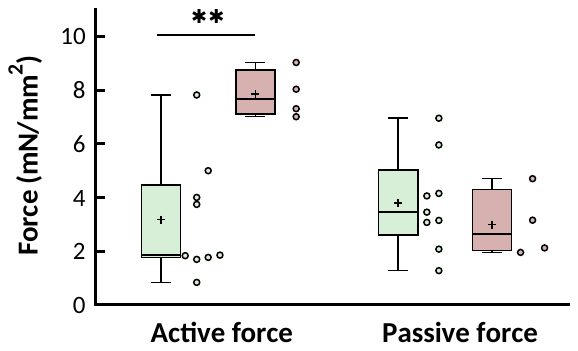

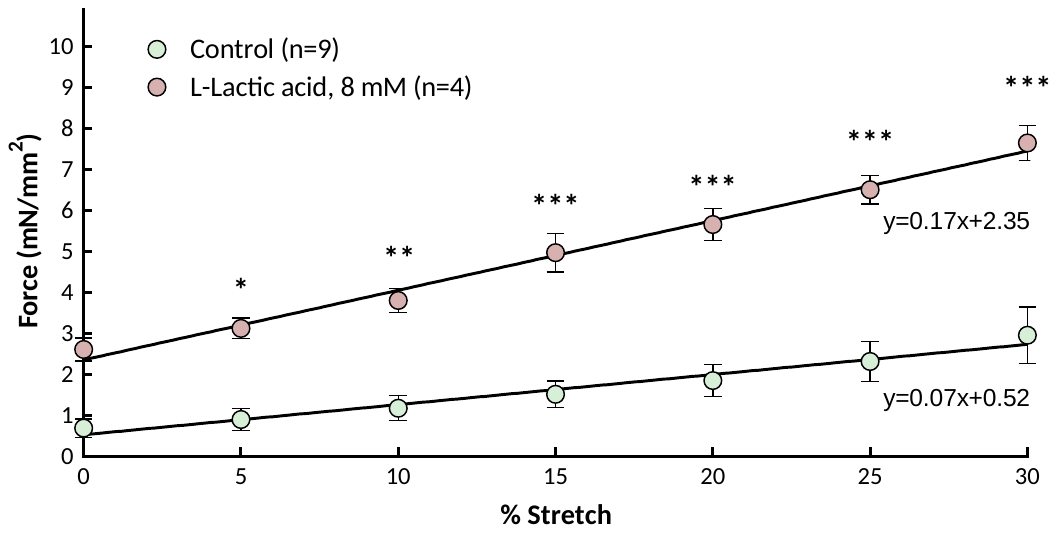

**A**

**B**

**C**

**TT	TT50%	TT90%**

**Supplementary Figure 5.** Effect of L-lactic acid on contractility of LMS from donor, healthy, human hearts. (A) Contraction force of LMS from human failing hearts under control conditions and after exposure to 8 mmol/L L-lactic acid for 48 hours. * (p<0.05), ** (p<0.01) and *** (p<0.001) indicate significant differences between both groups (two-way ANOVA followed by *post hoc* Bonferroni test, n=9 and 4 for control and 8 mM lactic acid-treated LMS, respectively). (B) Active and passive force of at maximum stretch (i.e., 30%). ** (p<0.01) indicates significant differences between both groups (Student’s t test). (C) Kinetics characteristics of LMS from human failing hearts at maximum contraction, including the time required to reach the peak amplitude of force (TTP, time to peak), and the time to decay from maximum force to 50% (TT50%, time to 50% decay) and to 90% (TT90%, time to 90% decay). Data for B and C are shown as box plot depicting median (horizontal line), mean (+), individual values (color symbols) and outlayers (x).

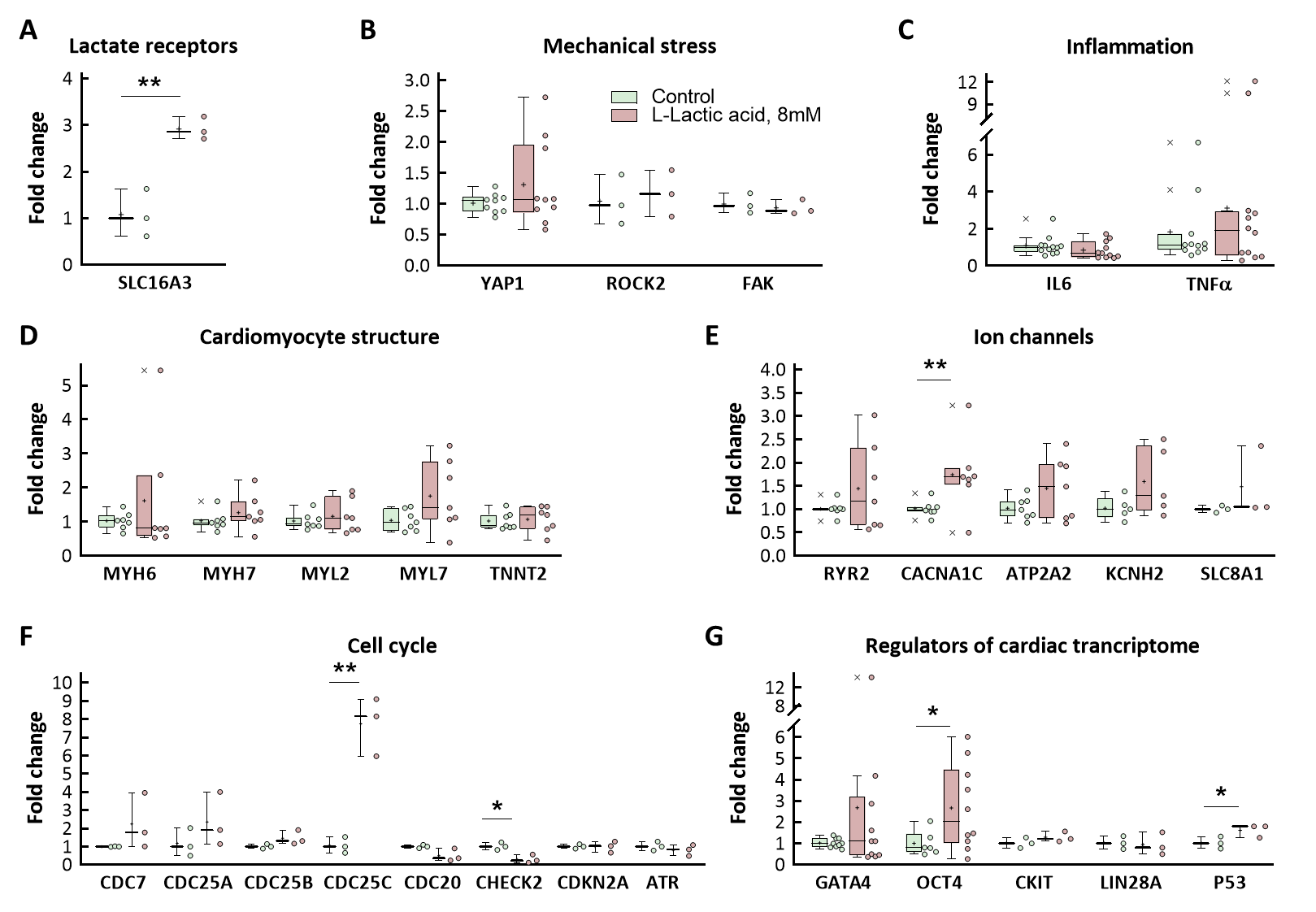

**Supplementary Figure 6.** Relative gene expression of LMS from human failing hearts exposed to control conditions or to incubation with 8 mmol/L L-Lactic acid for 48 hours. (A) Gene encoding for the lactate transporter MCT4. (B) Genes activated by mechanical stress. (C) Inflammatory genes. (D) Cardiomyocyte structural genes. (E) Genes involved in calcium handling. (F) Cell cycle genes. (G) Genes encoding for transcription factors and progenitor genes. * (p<0.05, Student’s t test) indicates significant differences between both groups (3 ≤ n ≤ 12). Data are shown as box plot depicting median (horizontal line), mean (+), individual values (color symbols) and outlayers (x).

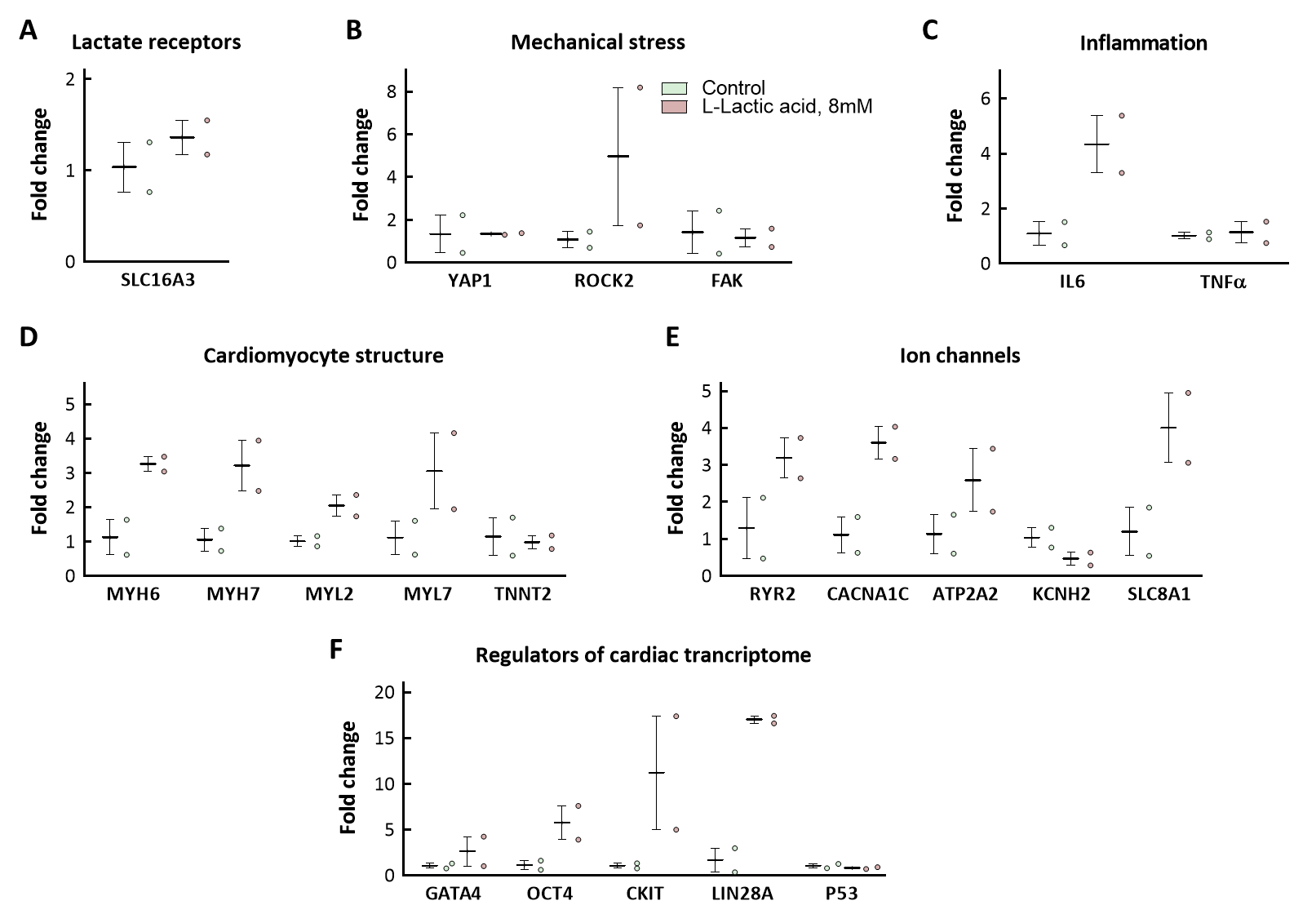

**Supplementary Figure 7 –** Relative gene expression of LMS from human healthy failing hearts exposed to control conditions or to incubation with 8 mmol/L L-Lactic acid for 48 hours. (A) Gene encoding for the lactate transporter MCT4. (B) Genes activated by mechanical stress. (C) Inflammatory genes. (D) Cardiomyocyte structural genes. (E) Genes involved in calcium handling. (F) Cell cycle genes. (G) Genes encoding for transcription factors and progenitor genes. Data are shown as box plot depicting median (horizontal line), mean (+), individual values (color symbols) and outlayers (x).

References.

1.	Valls-Lacalle L, Barba I, Miró-Casas E, Alburquerque-Béjar JJ, Ruiz-Meana M, Fuertes-Agudo M, Rodríguez-Sinovas A, García-Dorado D. Succinate dehydrogenase inhibition with malonate during reperfusion reduces infarct size by preventing mitochondrial permeability transition. *Cardiovasc Res* 2016;**109**:374–384.

2.	Li X, Yang Y, Zhang B, Lin X, Fu X, An Y, Zou Y, Wang JX, Wang Z, Yu T. Lactate metabolism in human health and disease. *Signal Transduct Target Ther* 2022;**7**.

3.	Watson SA, Scigliano M, Bardi I, Ascione R, Terracciano CM, Perbellini F. Preparation of viable adult ventricular myocardial slices from large and small mammals. *Nat Protoc* 2017;**12**:2623–2639.

4.	Watson SA, Duff J, Bardi I, Zabielska M, Atanur SS, Jabbour RJ, Simon A, Tomas A, Smolenski RT, Harding SE, Perbellini F, Terracciano CM. Biomimetic electromechanical stimulation to maintain adult myocardial slices in vitro. *Nat Commun* 2019;**10**.

5.	Ordoño J, Pérez-Amodio S, Ball K, Aguirre A, Engel E. The generation of a lactate-rich environment stimulates cell cycle progression and modulates gene expression on neonatal and hiPSC-derived cardiomyocytes. *Biomaterials advances* 2022;**139**.
